# Structural basis of ribosomal frameshifting during translation of the SARS-CoV-2 RNA genome

**DOI:** 10.1101/2020.10.26.355099

**Authors:** Pramod R. Bhatt, Alain Scaiola, Gary Loughran, Marc Leibundgut, Annika Kratzel, Angus McMillan, Kate M. O’ Connor, Jeffrey W. Bode, Volker Thiel, John F. Atkins, Nenad Ban

**Affiliations:** Department of Biology, Institute of Molecular Biology and Biophysics, ETH Zurich, Zurich, Switzerland; Schools of Biochemistry and Microbiology, University College Cork, Cork, Ireland, T12 XF62; Institute of Virology and Immunology, University of Bern, Bern, Switzerland; Laboratorium für Organische Chemie, Department of Chemistry and Applied Biosciences, ETH Zurich, Zurich, Switzerland.; Department of Infectious Diseases and Pathobiology, Vetsuisse Faculty, University of Bern, Bern, Switzerland; Department of Human Genetics, University of Utah, Salt Lake City, UT 84112-5330, USA; Graduate School for Cellular and Biomedical Sciences, University of Bern, Bern Switzerland

## Abstract

Programmed ribosomal frameshifting is the key event during translation of the SARS-CoV-2 RNA genome allowing synthesis of the viral RNA-dependent RNA polymerase and downstream viral proteins. Here we present the cryo-EM structure of the mammalian ribosome in the process of translating viral RNA paused in a conformation primed for frameshifting. We observe that the viral RNA adopts a pseudoknot structure lodged at the mRNA entry channel of the ribosome to generate tension in the mRNA that leads to frameshifting. The nascent viral polyprotein that is being synthesized by the ribosome paused at the frameshifting site forms distinct interactions with the ribosomal polypeptide exit tunnel. We use biochemical experiments to validate our structural observations and to reveal mechanistic and regulatory features that influence the frameshifting efficiency. Finally, a compound previously shown to reduce frameshifting is able to inhibit SARS-CoV-2 replication in infected cells, establishing coronavirus frameshifting as target for antiviral intervention.

## Main Text

Ribosomal frameshifting, a process during which the reading frame of translation is changed in the middle of the coding sequence, is one of the key events during translation of the severe acute respiratory syndrome coronavirus 2 (SARS-CoV-2) positive sense single-stranded RNA genome. This so-called programmed -1 translational frameshifting is conserved in all coronaviruses and is necessary for synthesis of viral RNA dependent RNA polymerase (Nsp12 or RdRp) and downstream viral non-structural proteins encoding core enzymatic functions involved in capping of viral RNA, RNA modification and processing, and RNA proof-reading. Although the translational machinery typically prevents frameshifting as a potential source of one of the most disruptive errors in translation (*1*, *2*), many viruses rely on programmed ribosomal frameshifting to expand and fine-tune the repertoire and stoichiometry of expressed proteins(*3*).

Programmed -1 frameshifting in SARS-related coronaviruses occurs at the slippery sequence U_UUA_AAC in the context of a 3’ stimulatory RNA sequence that was predicted to form a 3-stemmed pseudoknot structure (*4*), the importance of which was tested by our lab and others (*5*, *6*). The frameshifting occurs with high efficiency (25-75% depending on the system used (*5*–*9*) and changes the reading frame to UUU_AAA_C (*10*) (Fig. 1A). A putative secondary structure element in the viral RNA that forms a loop upstream of the shift site has been proposed to play an attenuating role in frameshifting and is referred to as the 5’ attenuator loop (*11*). Maintaining the precise level of coronavirus frameshifting efficiency is crucial for viral infectivity, evidenced by the remarkable fact that mutation of a single nucleotide in the frameshifting region of the SARS-CoV RNA results in a concomitant abrogation of viral replication (*12*). Therefore, the criticality of pseudoknot-dependent -1 ribosomal frameshifting in the propagation of SARS-related coronaviruses, a process that does not occur in human cells, presents itself as an opportune drug-target with minimal tolerance for drug-resistant mutations.

**Fig. 1.**
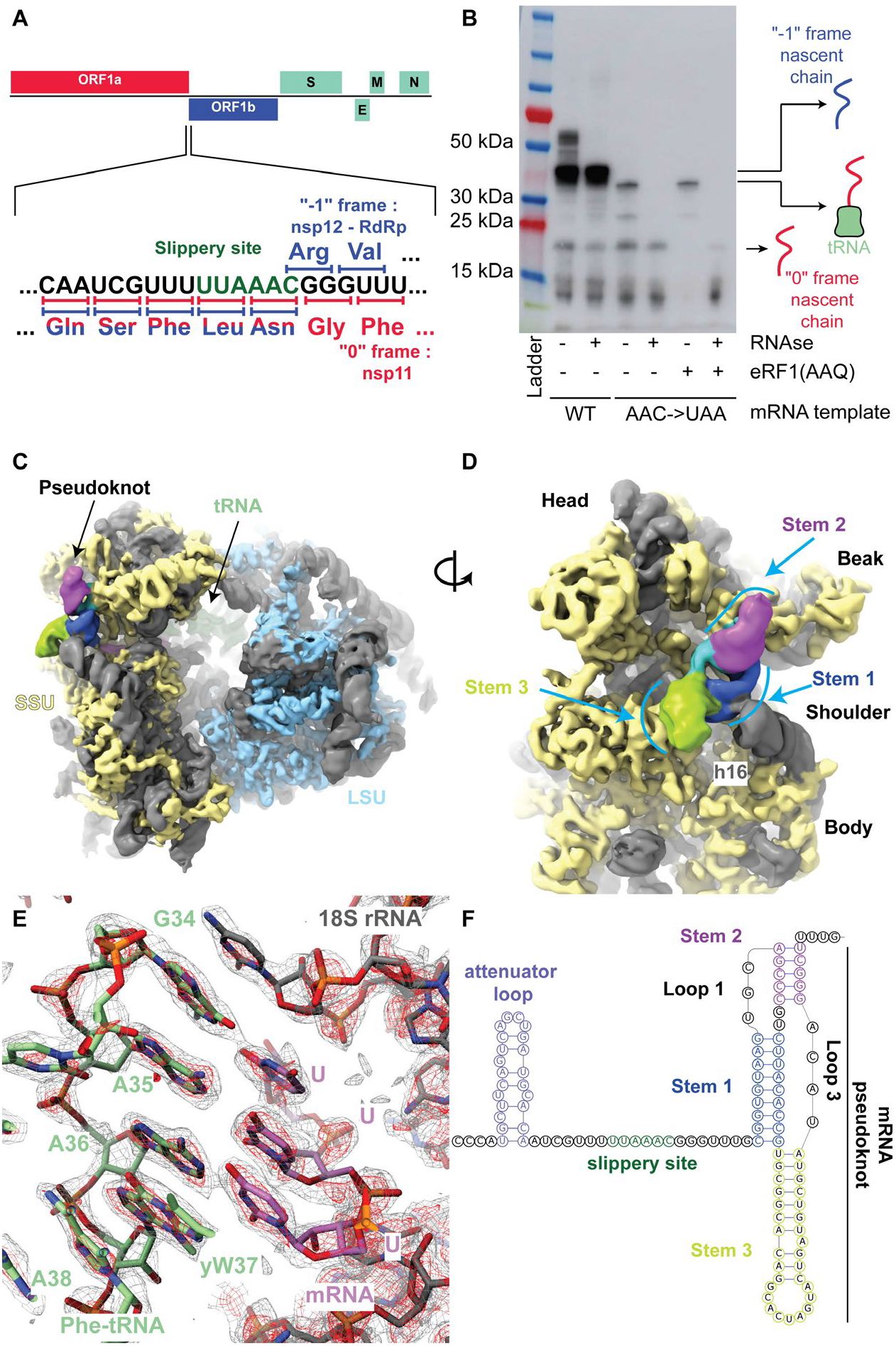
The SARS-Cov-2 pseudoknot interacts with the ribosome and pauses translation upstream of the slippery site. (**A**) Schematic of the SARS-CoV-2 main ORF. In the close up view of the frameshift event, codons and corresponding amino acids are shown. During -1 frameshifting, the ‘slippery site’ codons UUA (Leu) and AAC (Asn) are the last codons decoded in the 0 frame. Upon -1 frameshifting of the AAC codon to AAA, translation resumes at the CGG (Arg) triplet, where elongation proceeds uninterrupted to produce full-length Nsp12. (**B**) In vitro translation reaction depicting pausing at the frameshift site. Efficient frameshifting is observed for the WT template. Samples for cryo-EM originally intended to be trapped by dominant negative eRF1 (AAQ) show a tRNA-bound pause in proximity of the frameshift site. The tRNA-associated band is lost upon RNAse treatment. Reactions without added eRF1 (AAQ) produce a similarly paused product. (**C**) Overview of the density low pass filtered to 6Å with the pseudoknot found close to the entry of the mRNA channel on the small subunit (SSU). The SSU proteins are colored in yellow, the large subunit (LSU) proteins in blue and the rRNA in grey. The pseudoknot is colored according to its secondary structure as in panel (F), and the P-site tRNA is colored in dark green. (**D**) Close-up view of the pseudoknot from the solvent-exposed side of the SSU. Helix h16 of the 18S rRNA interacts with the base of Stem 1. Unpaired loop-forming nucleotides are colored in cyan. (**E**) P-site codon-anticodon interactions reveal a Phe (UUU) codon interacting with Phe-tRNA. (**F**) Schematic of the revised secondary structure elements in the pseudoknot necessary for -1 PRF with different functional regions labeled and colored accordingly.

Due to its importance in the life cycle of many important viruses and coronaviruses in particular, programmed frameshifting has been extensively studied using a range of structural and functional approaches (*3*). The structure of a 3’ stimulatory pseudoknot in isolation or in context of the viral genome has been proposed recently by various groups using techniques that include molecular dynamics, nuclease mapping, in vivo SHAPE, NMR and cryo-EM (*6*, *13*–*16*). Furthermore, a ribosomal complex with a frameshift stimulatory pseudoknot from the avian infectious bronchitis virus was reported at low resolution (*17*). Here, to provide a structural and mechanistic description of the events that ensue during ribosomal frameshifting, we investigated mammalian ribosomes captured in distinct functional states during translation of a region of SARS-CoV-2 genomic RNA where -1 programmed frameshifting occurs.

### Structure determination of a frameshifting-primed ribosomal complex

We initially intended to capture a 0 frame, pre-frameshift ribosomal complex by first introducing a stop codon in place of the second codon of the slippery site (U_UUA_AAC to U_UUA_UAA), achieving stalling by means of utilizing a mutant eRF1 (AAQ) that is unable to release the nascent polypeptide and would, therefore, trap the ribosome in the act of decoding the mutant frameshift site in the 0 frame. Translating complexes were prepared in an *in vitro* translation reaction using a rabbit reticulocyte lysate (RRL) system generated in-house (see Methods). The ribosomes were programmed with mRNA harboring a region of the SARS-CoV-2 genome that encodes proteins Nsp10 (C-terminus), Nsp11 and the majority of Nsp12. Western blotting showed that when using the WT RNA template, frameshifting was efficient, while the stop codon mutation prevented frameshifting and led to ribosome stalling when eRF1 (AAQ) was present (Fig. 1B). Curiously, we also observed a prominent band corresponding to a tRNA-bound product paused in the vicinity of the frameshift site even in the absence of eRF1 (AAQ).

Indeed, cryo-electron microscopic 3D reconstruction of RNCs purified from the reactions supplemented with eRF1 (AAQ) revealed that the majority of ribosomal particles corresponded to a paused ribosome with no detectable eRF1 (AAQ) (fig. S1). The 80S ribosomes were found captured in the process of translation with P- and E- site tRNAs bound, and with a nascent polypeptide chain in the ribosomal tunnel.

Incidentally, the 2.2 Å resolution obtained for our reconstruction allowed us to build the most accurate structure of a mammalian 80S ribosome so far and directly visualize virtually all rRNA modifications proposed for the human ribosome based on quantitative mass spectrometry, which is not surprising since all modified residues are also conserved in rabbit rRNAs (*18*) (fig. S2 and S3). Notably, these results are not consistent with the modifications of the human rRNAs recently proposed based on interpretations of EM maps (*19*).

Importantly, further classification revealed a prominent density for a complete 3’ frameshifting stimulatory pseudoknot at the entry of the mRNA channel on the 40S subunit (Fig. 1C–D). The resolution of this reconstruction ranged from 2.3 Å at the core of the ribosome to ~7 Å at the periphery, where the most flexible regions of the pseudoknot are located (fig. S1 and S2). Based on the high-resolution maps that allowed visualization of the codon-anticodon interactions and modifications in the tRNA (Fig. 1E, fig. S4A-B), we could unequivocally determine that a Phe-tRNA(Phe) was bound at the P-site (*20*). This implied that the ribosome is paused by the downstream pseudoknot such that the P-site tRNA interacts with the UUU codon just prior to the first codon of the frameshift site (Fig. 2A).

**Fig. 2.**
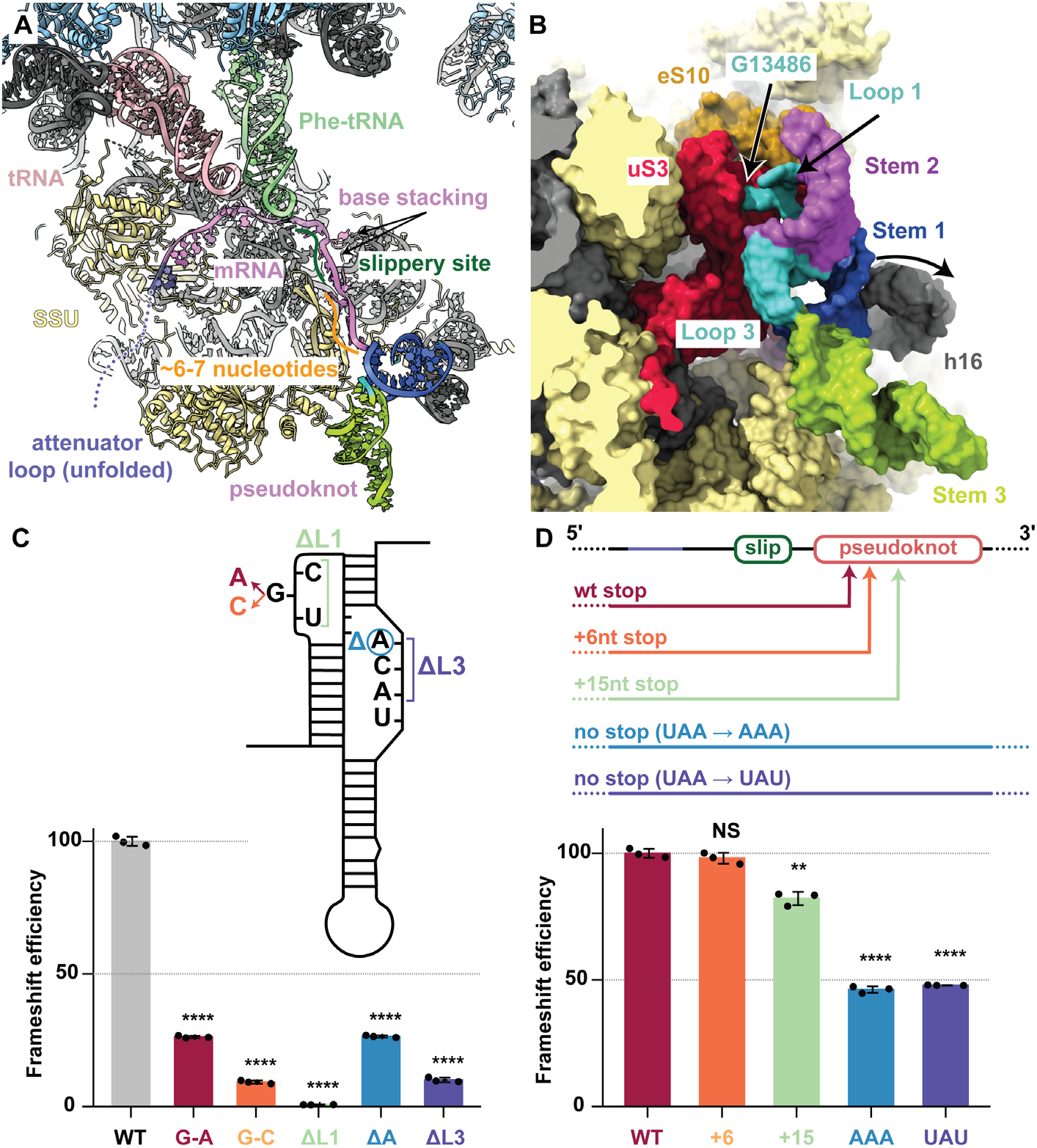
Critical features of the ribosome-bound pseudoknot. (**A**) Overview of the frameshift-primed state. The stimulatory pseudoknot pauses the ribosome at the penultimate codon (UUU) of the slippery site, with P- (green) and E- (pink) sites occupied by tRNAs, and an empty A-site awaiting decoding in the non-rotated state. The length of the spacer region (orange) is critical for exact positioning of the pseudoknot as the spacer exerts tension at the entry of the mRNA channel. (**B**) The backbone of Loop 1 (UGC) (cyan) of the pseudoknot interacts with the N-terminal domain of uS3 (red) and the C-terminal tail of eS10 (orange). mRNA residue G13486 is flipped out and interacts with uS3. (**C**) Mutagenesis experiments using dual luciferase assays in HEK cells indicate that the G13486 interaction is specific. Mutation of G13486 to other residues leads to a marked reduction in frameshifting efficiency, and deletion of Loop 1 (Δ L1) completely abolishes frameshifting. Similarly, deletion of a single nucleotide (A13537) in Loop 2 reduces frameshifting, while deletion of the entire loop (ΔL2) abolishes frameshifting. Normalized (Firefly/Renilla) luciferase activites were calculated for each construct as a percentage of their individual normalized in-frame controls. Data are presented as mean values ± standard deviations of three biological replicates (sets of translation reactions) averaged after three measurements, with error bars representing standard deviations. (**D**) Mutagenesis experiments using dual luciferase reporter assays in HEK cells show that the position of the 0 frame stop codon influences frameshifting. Leaving the pseudoknot unaltered, incremental increase in the distance of the 0 frame stop codon from the frameshift site leads to a concomitant decrease in frameshifting levels. Loss of the stop codon in 0 frame leads to a sharp decline in frameshifting levels.

### The SARS-Co-V-2 RNA pseudoknot specifically interacts with ribosomal proteins and 18S rRNA

The intermediate local resolution (5-7 Å) of the cryo-EM map in the area of the pseudoknot allowed us to visualize the overall fold of the RNA and readjust its previously predicted secondary structure (*13*–*16*, *21*) (Fig. 1E). The stimulatory pseudoknot forms an H-type pseudoknot with Stem 1 and Stem 2 coaxially stacked on top of each other to form a quasi-continuous helix, while Stem 3 stands out almost perpendicular to this plane (Fig. 1D and 2B). This corkscrew-like formation provides a bulky and well-structured obstacle wedged at the mRNA entry channel, having the potential to resist unwinding by the helicase activity of the ribosome and generating tension on the upstream mRNA up to the decoding center. Stem 1 of the pseudoknot forms a 9 bp helix which is GC rich at the bottom (Fig. 1F). The penultimate nucleotides of the ‘spacer region’ prior to Stem 1 are located at the mRNA entry tunnel, where they interact with several basic residues in the C-terminal domain of uS3 on one side and are supported by uS5 from the other, with an additional weak contact being made from the C-terminal end of eS30. uS3 and eS30 are primary components of the ribosome helicase and uS5 has been proposed to be a component of the ribosomal helicase processivity clamp at the mRNA entry site (*22*, *23*). The observed distance between the P-site UUU codon and Stem 1 of the pseudoknot underscore the critical dependence of the frameshifting efficiency on the length of the spacer region (*24*). Translocation to the next codon would place the frameshifting codon UUA into the P-site, with a simultaneous increase in the tension of the mRNA and unwinding of the GC-rich base of Stem 1 upon entering into the mRNA entry channel.

The pseudoknot structure also reveals a hitherto unobserved and possibly unappreciated role for the distal site of the mRNA entrance channel in helicase activity. While mRNA unwinding studies outside the mRNA entrance channel have so far implicated only a helix in the C-terminal domain of uS3 (*25*), we notice that Loop 1 of the pseudoknot contacts the N-terminal domain of uS3 as well as the C-terminal tail of eS10 (Fig. 2B and fig. S4D), whereas the flipped-out base G13486 in this loop forms specific interactions (Fig. 2B). Furthermore, as the pseudoknot is located at the entry to the mRNA channel, helix h16 of the 18S rRNA is noticeably pushed outwards due to a direct contact with the minor groove of Stem 1 (Fig. 2B, fig. S5A). Since the pseudoknot wedges between the head and the body of the small ribosomal subunit, it would restrict their relative motions that need to take place during translocation. This is consistent with the studies on dynamics of coronavirus frameshifting, which revealed that the mechanism of -1 frameshifting involves restriction of small subunit head motion (*26*).

The structure also reveals another key aspect of the architecture of the pseudoknot as the ribosome encounters it. The start of the pseudoknot is shifted relative to the predicted secondary structure (*13*–*16*, *21*) by two nucleotides. The two opposed nucleotides, which were assumed to base pair with Stem 1, are actually forming the start of Stem 3 by pairing with bases predicted to be in the single-stranded linker 2 (Fig. 1F, fig. S5B-C). Our cryo-EM density reveals that Loop 3 accommodates a total of 4 nucleotides, three of which were originally attributed to Stem 2. Thus, we observe that Loop 3 is shifted and expanded relative to the initially predicted secondary structures (*13*–*16*, *21*).

To functionally support our structural findings and confirm the nature and specificity of the pseudoknot interactions, we performed structure-guided mutagenesis experiments using dual luciferase reporter assays in HEK cells (see Methods) and monitored the frameshifting efficiency relative to the WT (Fig. 2C). Mutation of G13486 of Loop 1 to another purine reduced the frameshifting efficiency to 30% of the WT level, and mutation of this base to a pyrimidine further reduced frameshifting to 15%. As expected from our structural data, deletions of the nucleotides of the spacer regions also had a deteriorating effect on frameshifting. Loss of Loop 1 entirely abolished frameshifting. Deletion of a single nucleotide of Loop 3, which would not be expected to have an effect on the folding of the pseudoknot based on the previously suggested secondary structures where it is flipped out (*21*), diminished the frameshifting rate to 25% of the WT level. Loss of the entire Loop 3 reduced frameshifting to 10% of WT levels.

### Frameshifting efficiency depends on the position of the “0” frame stop codon

In SARS-CoV-2, the 0 frame stop codon is located 5 codons downstream of the frameshift site and is a constituent of Stem 1. The placement of the stop codon in such proximity to the frameshift site is a common feature in coronaviruses, and its presence in a critical region of the stimulatory pseudoknot prompted us to probe the effect of the distance of the 0 frame stop codon on frameshifting. To this end, knowledge of the 3D structure of the pseudoknot helped us to confidently manipulate the stop codon without hampering pseudoknot formation. We introduced mutations to incrementally extend the stop codon from the WT position and to completely remove the occurrence of a stop codon in the 0 frame (Fig. 2D). While introducing a stop codon 6 nucleotides downstream of the WT position only marginally decreased the frameshifting rate (98% of WT), a stronger attenuation was observed when the distance of the stop codon was increased to 15 nucleotides from the WT stop (80% of WT). Finally, removal of the stop codon by two different point mutations led to a reduction of frameshifting efficiency to 45% of WT levels.

Taken together, these observations suggest that the stop codon position plays an important role in maintaining optimum frameshift efficiency. We propose that the stop codon serves to prevent the closely trailing ribosome from encountering an RNA that was unfolded by the leading ribosome. In this case, upon encountering a stop codon, termination and subunit disassembly will occur, which will provide an opportunity for the pseudoknot to refold without the constrains of the mRNA channel (see Conclusions). This mechanism, consistent with our biochemical data, increases the efficiency of frameshifting to the levels required by SARS-CoV-2 and may be used by viruses in general when high-efficiency frameshifting is required.

### Inhibition of viral replication by a compound that targets the SARS-CoV-2 pseudoknot

The extreme sensitivity of the coronavirus to the finely controlled frameshifting levels (*12*) may present an opportunity to develop compounds that interfere with the frameshifting process and thus inhibit replication of the virus. Using computational modeling and reporter assays, compounds that have been predicted to bind the pseudoknot and inhibit SARS-CoV-2 frameshifting were described (*21*, *27*), but never tested with respect to their ability to inhibit viral replication. To demonstrate that the inhibition of frameshifting is a plausible strategy for drug development, we synthesized MTDB, one of these previously described compounds (*21*, *27*, *28*), and tested whether it is able to reduce viral levels in infected African green monkey Vero cells (Fig. 3, see Methods). The compound showed no cellular toxicity and resulted in a 3 orders of magnitude reduction of SARS-CoV-2 titer, with the half maximal inhibitory concentration (IC50) of 48 μM (Fig. 3). Although this potency range is still far from what would be expected from a potential drug candidate, it nevertheless provides a starting point for high-throughput screening, and establishes that frameshifting is a viable target for therapeutic intervention against SARS-CoV-2.

**Fig. 3.**
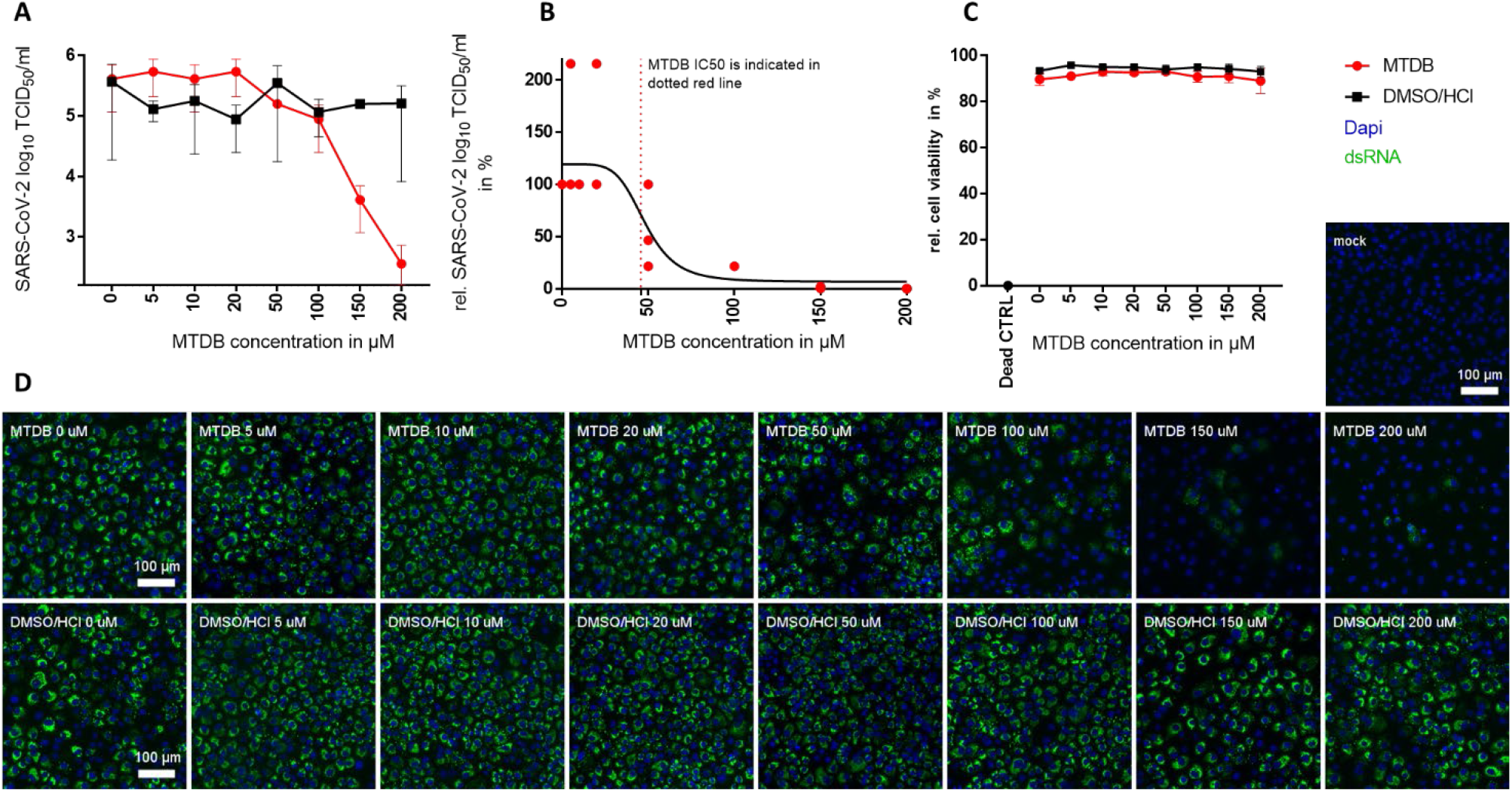
Effect of MTDB on SARS-CoV-2 (MOI 0.01) replication in VeroE6 cell lines. Vero E6 cells were infected with SARS-CoV-2 and treated with MTDB at indicated concentrations. (**A**) SARS-CoV-2 titers are displayed in TCID50/ml in a log10 scale, 24 hours post infection. (**B**) Relative SARS-CoV-2 titers are depicted in percent and non-linear regression (inhibitor vs. response) was performed using Graph Pad Prism 7.04 to determine MTDB IC50 of 48 μM. MTDB IC50 is indicated as red dotted line. (**C**) The relative VeroE6 cell viability displayed in percent. The MTDB-treated condition is designated in red, the solvent control in black. Dilutions of MTDB range from 0-200 μM, respective volumes of 20% DMSO/100 mM HCL were used as solvent control. A mean of three independent experiments with SD (error bars) is shown. (**D**) Representative immunofluorescence images of one out of three replicates are shown, scale bar is 100 μm, SARS-CoV-2 double stranded RNA is shown in green, DAPI is shown in blue.

### The nascent chain modulates the frameshift efficiency in SARS-CoV-2

Strikingly, in the reconstruction of the paused translating ribosome, the nascent chain that corresponds to the viral polyprotein was visible along the entire length of the ribosomal exit tunnel (Fig. 4A). The density corresponded to the C-terminal region of Nsp10, which is the activator of the viral proofreading exonuclease and N7-methyltransferase Nsp14 (*29*, *30*) and then, depending on the frameshifting event, continues as either the viral RNA-dependent RNA polymerase Nsp12 (*5*) or protein Nsp11, the function of which is yet unknown (Fig. 1A and 4B). The nascent chain makes several specific interactions with the ribosomal tunnel, one of which is at the constriction site where arginine 4387 of Nsp10 interacts with A1555 of the 28S rRNA (corresponding to A1600 in humans, numbering according to PDB 6EK0 (*19*)) and is stabilized by the preceding leucine 4386 (Fig. 4C). Interestingly, these two amino acids are very well conserved across multiple coronaviruses (fig. 4G), although they are located in the unstructured C-terminal region of Nsp10 and therefore considered not to be important for the fold of the protein (*31*).

**Fig. 4.**
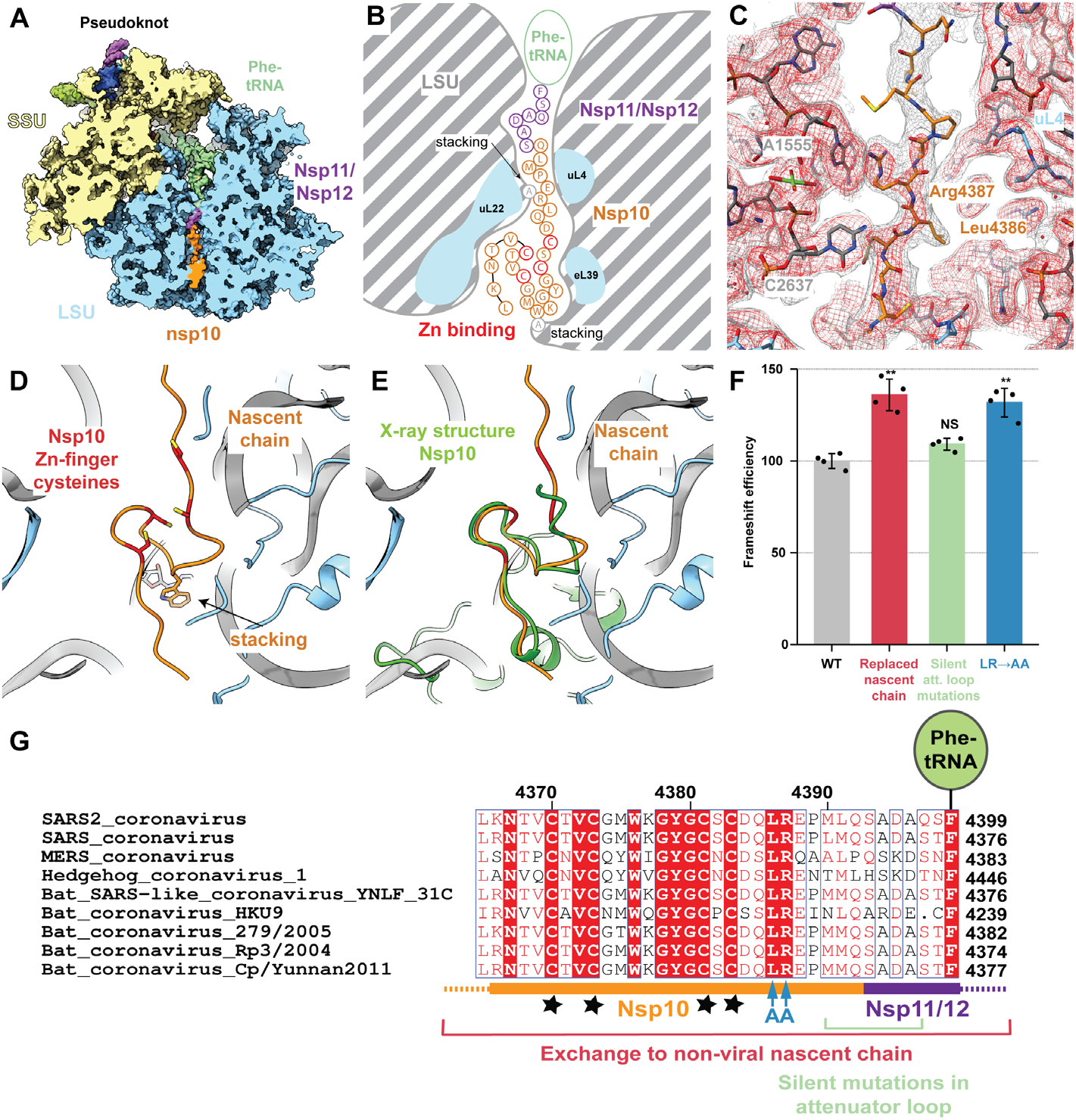
The nascent viral polypeptide co-translationally folds and specifically interacts with the ribosomal tunnel. (**A**) Cross-section of the pseudoknot-paused ribosome structure showing the exit tunnel. The nascent C-terminus of Nsp10 (orange) and the N-terminus of Nsp11/12 (purple) are visible from the PTC to the periphery of the ribosome exit tunnel (LSU in blue). (**B**) Schematic representation of the path of nascent peptide along the exit tunnel. Arg 4387 stacks with 28S rRNA residue A1555 at the constriction site. Further down, where the tunnel widens, the C-terminal zinc finger domain of Nsp10 folds co-translationally, with Trp 4376 stacking on A2261 of 28S rRNA. (**C**) Well-ordered density is visible for Arg 4387 of Nsp10 as it stacks onto A1555 of 28S rRNA at the constriction site and is stabilized by Leu 4386. The structure is shown within the cryo-EM map contoured at two different levels (grey and red). (**D**) and (**E**) The overlay of the co-translationally folded zinc finger domain with the crystal structure of Nsp10 (green, PDB 2FYG (31)) reveals the structural similarity. (**F**) Probing the role of nascent chain interactions with the ribosome exit tunnel using an RRL in vitro system. Mutations of the interacting residues were tested for their effect on frameshifting. Replacement of the entire nascent chain with an unrelated sequence leads to a 35% increase in frameshifting, which is only in part due to loss of the 5’ attenuator loop. Interactions around the constriction site likely serve to attenuate frameshifting, as replacement of the interacting Arg 4387 and stabilizing Leu 4386 with Ala increases frameshifting by 30%. (**G**) Alignment of SARS2 with closely related sequences of other coronaviruses highlighting the conservation of the mutated residues (colored as in (F)). The shown sequence stretch encompasses the C-terminal zinc finger domain of Nsp10 (orange) and parts of Nsp11/Nsp12 (purple) visible in our reconstruction. Nascent chain residues Leu 4386 and Arg 4387 interacting with the ribosomal exit tunnel are strictly conserved, while the conservation of neighboring residues is lower. Stars represent the four cysteines of the Nsp10 zinc finger.

Further down the tunnel, the C-terminal end of Nsp10 adopts a partially-folded zinc finger motif (Fig. 4D–E), which upon superposition reveals similarity with the corresponding fully folded C-terminal domain previously observed in the crystal structure of SARS-CoV Nsp10 (*31*). Tryptophan 4376 located between the two pairs of cysteines that form the zinc finger stacks with A2261 (A2418), an interaction that might serve to promote the change of nascent chain direction and facilitate folding of the zinc finger at the end of the exit tunnel. Co-translational events, such as insertion of a transmembrane domain at the exit of the ribosomal tunnel, was shown to promote -1 ribosomal frameshifting in alphaviruses (*32*).

To investigate whether the observed contacts between the nascent chain and the ribosomal tunnel are specific, and whether these interactions and co-translational folding of Nsp10 might play a role in modulating the frameshifting process, we employed our dual luciferase reporter assay to measure the frameshifting efficiency of WT and mutant nascent chain sequence constructs. As our measurements in HEK cells did not reveal an appreciable change of frameshift efficiency, we carried out the same experiments *in vitro* using RRL to monitor the effects in a single mRNA setup. Replacement of the entire nascent chain with an unrelated sequence leads to a 35% increase in frameshifting (Fig. 4F). The extent of this change is clearly not due to loss of the 5’ attenuator loop, as indicated by an experiment where silent attenuator loop mutations result in only a slight increase in frameshifting. Mutation of the leucine 4386 and arginine 4387 to alanine led to a considerable (30%) increase in frameshifting (Fig. 4F–G), implying that these nascent chain interactions with the ribosomal exit tunnel play an important role in regulating frameshifting levels, possibly mechanistically akin to the well-studied SecM stalling system in bacteria (*33*), where it was shown that co-translational folding and the translocon-induced mechanical force can rescue the stall induced by interactions between the nascent chain and the ribosomal tunnel (*34*). These observations also imply that any cellular nascent-chain factors (*35*, *36*), or even the intracellular concentration of Zn^2+^, which is known to change in response to viral infections (*37*), might influence the rate of frameshifting.

## Conclusions

Our results provide a mechanistic description of frameshifting that occurs during translation of the SARS-CoV-2 genome and reveal the features that may be exploited by the virus to finely control the stoichiometry of viral proteins at different stages of infection (Fig. 5). Interfering with the frameshifting process at the level of nascent chain interactions with the ribosomal tunnel, at the level of RNA folding that leads to the formation of the frameshift stimulatory pseudoknot, or to perturb the interactions between the pseudoknot and the mRNA channel, represent a viable strategy in our search for new drugs against SARS-CoV-2, the virus that is currently causing the global COVID-19 pandemic. Our results will also be useful for understanding the mechanism of programmed ribosomal “-1” frameshifting employed by many other medically important viruses.

**Fig. 5.**
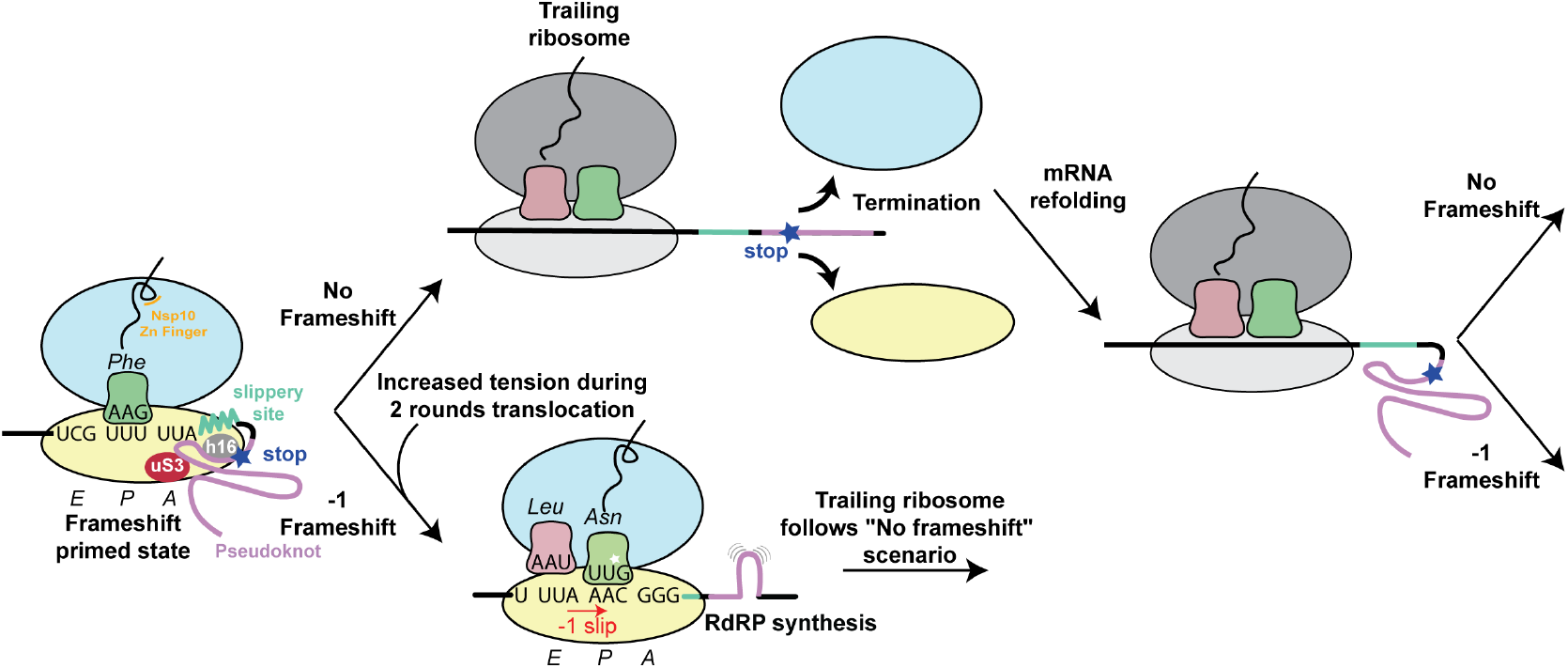
Structure based model for -1 programmed frameshifting in coronaviruses and its regulation. The observed interactions between the pseudoknot and the ribosome prime the system for frameshifting. The features of the pseudoknot and the interactions between the nascent chain and the ribosomal tunnel play a role in the efficiency of frameshifting. The efficiency of frameshifting is increased by the presence of a stop codon near the frameshifting site. Ribosomes that progress beyond the frameshifting site in the 0 frame quickly terminate and disassemble, thereby increasing the chances that the pseudoknot will refold before it is encountered by the closely trailing ribosome. The trailing ribosome in turn encounters the pseudoknot, which increases the possibility of undergoing -1 frameshifting.

## Supporting information

Materials and Methods and all supplementary material

## Acknowledgments

We thank A. Jomaa for advice on cryo-EM data processing and A. Picenoni for help with the grid preparation. We are indebted to the ETH scientific center for optical and electron microscopy (ScopeM) for access to electron microscopes, and in particular to M. Peterek and D. Boehringer.

## Funding

This work was supported by the Swiss National Science Foundation (SNSF) to NB and VT (grant 31003A_182341 and 310030_173085) via the National Center of Excellence in RNA and Disease (project funding 138262) to N. Ban and V. Thiel ; The Irish Research Council Advanced Laureate (IRCLA/2019/74) to J.F.A., who thanks Prof. G.F. Fitzgerald for funds from Carbery Group Ltd.

## Author contributions

PRB and NB initiated the project, and together with JFA designed the experiments. PRB carried out biochemical experiments and sample preparation. AS and PRB prepared grids. AS carried out data collection and calculated EM maps. AS and ML performed molecular model building and refinement. AS, ML, PRB and NB interpreted the structure. PRB, AS, ML and NB drafted the manuscript. AS prepared the figures. GL and KMOC performed dual-luciferase reporter assays. AM prepared and purified the drug-like ligand MTDB. AK and VT formulated experiments to test viral infected cells. AK carried out testing of ligand on viral-infected cells. All authors contributed to the final version of the manuscript

## Competing interests

The authors declare no competing interest.

## Data and materials availability

The structure and cryo-EM map for the high-resolution reconstruction are available in the Protein Data Bank as PDB ID XXXX and in the Electron Microscopy Data Bank as EMD-YYYY, respectively. The structure refined into the further classified particle set reconstruction and the corresponding maps are available as PDB-ZZZZ and EMD-WWWW, respectively.

## Supplementary Materials

Materials and Methods

Figures S1-S5

Tables S1-S2

## Notes

### Competing Interest Statement

The authors have declared no competing interest.

## References

1. J. Parker, Errors and alternatives in reading the universal genetic code. Microbiol. Rev. 53, 273–298 (1989).

2. J. M. Ogle, A. P. Carter, V. Ramakrishnan, Insights into the decoding mechanism from recent ribosome structures. Trends Biochem. Sci. 28, 259–266 (2003).

3. J. F. Atkins, G. Loughran, P. R. Bhatt, A. E. Firth, P. V. Baranov, Ribosomal frameshifting and transcriptional slippage: From genetic steganography and cryptography to adventitious use. Nucleic Acids Res. 44 (2016), doi:10.1093/nar/gkw530.

4. F. Dos Ramos, M. Carrasco, T. Doyle, I. Brierley, Programmed -1 ribosomal frameshifting in the SARS coronavirus. Biochem. Soc. Trans. 32, 1081–1083 (2004).

5. P. V. Baranov, C. M. Henderson, C. B. Anderson, R. F. Gesteland, J. F. Atkins, M. T. Howard, Programmed ribosomal frameshifting in decoding the SARS-CoV genome. Virology. 332, 498–510 (2005).

6. E. P. Plant, G. C. Pérez-Alvarado, J. L. Jacobs, B. Mukhopadhyay, M. Hennig, J. D. Dinman, A three-stemmed mRNA pseudoknot in the SARS coronavirus frameshift signal. PLoS Biol. 3, 1012–1023 (2005).

7. I. Brierley, P. Digard, S. C. Inglis, Characterization of an efficient coronavirus ribosomal frameshifting signal: Requirement for an RNA pseudoknot. Cell. 57, 537–547 (1989).

8. N. Irigoyen, A. E. Firth, J. D. Jones, B. Y. W. Chung, S. G. Siddell, I. Brierley, High-Resolution Analysis of Coronavirus Gene Expression by RNA Sequencing and Ribosome Profiling. PLoS Pathog. 12 (2016), doi:10.1371/journal.ppat.1005473.

9. Y. Finkel, O. Mizrahi, A. Nachshon, S. Weingarten-Gabbay, D. Morgenstern, Y. Yahalom-Ronen, H. Tamir, H. Achdout, D. Stein, O. Israeli, A. Beth-Din, S. Melamed, S. Weiss, T. Israely, N. Paran, M. Schwartz, N. Stern-Ginossar, The coding capacity of SARS-CoV-2. Nature (2020), doi:10.1038/s41586-020-2739-1.

10. V. Thiel, K. A. Ivanov, Á. Putics, T. Hertzig, B. Schelle, S. Bayer, B. Weißbrich, E. J. Snijder, H. Rabenau, H. W. Doerr, A. E. Gorbalenya, J. Ziebuhr, Mechanisms and enzymes involved in SARS coronavirus genome expression. J. Gen. Virol. 84 (2003), pp. 2305–2315.

11. M. C. Su, C. Te Chang, C. H. Chu, C. H. Tsai, K. Y. Chang, An atypical RNA pseudoknot stimulator and an upstream attenuation signal for -1 ribosomal frameshifting of SARS coronavirus. Nucleic Acids Res. 33, 4265–4275 (2005).

12. E. P. Plant, R. Rakauskaite, D. R. Taylor, J. D. Dinman, Achieving a golden mean: mechanisms by which coronaviruses ensure synthesis of the correct stoichiometric ratios of viral proteins. J. Virol. 84, 4330–4340 (2010).

13. R. Rangan, I. N. Zheludev, R. J. Hagey, E. A. Pham, H. K. Wayment-Steele, J. S. Glenn, R. Das, RNA genome conservation and secondary structure in SARS-CoV-2 and SARS-related viruses: A first look. RNA. 26, 937–959 (2020).

14. K. Zhang, I. N. Zheludev, R. J. Hagey, M. Teng, P. Wu, R. Haslecker, Y. J. Hou, R. Kretsch, G. D. Pintilie, R. Rangan, W. Kladwang, S. Li, E. A. Pham, C. Bernardin-Souibgui, R. S. Baric, T. P. Sheahan, V. D′souza, J. S. Glenn, W. Chiu, R. Das, Cryo-electron microscopy and exploratory antisense targeting of the 28-kDa Frameshift Stimulation Element from the SARS-CoV-2 RNA genome. bioRxiv. (2020) doi:10.1101/2020.07.18.209270.

15. T.C.T. Lan, M. F. Allan, L. E. Malsick, S. Khandwala, S. S. Y. Nyeo, M. Bathe, A. Griffiths, S. Rouskin, Structure of the full SARS-CoV-2 RNA genome in infected cells. bioRxiv. (2020) doi: 10.1101/2020.06.29.178343L.

16. N. C. Huston, H. Wan, R. De Cesaris, A. Tavares, C. Wilen, A. Marie, N. C. Huston, H. Wan, C. Huston, Comprehensive in-vivo secondary structure of the SARS-CoV-2 genome reveals novel regulatory motifs and mechanisms. bioRxiv. (2020) doi: 10.1101/2020.07.10.197079

17. O. Namy, S. J. Moran, D. I. Stuart, R. J. C. Gilbert, I. Brierley, A mechanical explanation of RNA pseudoknot function in programmed ribosomal frameshifting. Nature. 441, 244–247 (2006).

18. M. Taoka, Y. Nobe, Y. Yamaki, K. Sato, H. Ishikawa, K. Izumikawa, Y. Yamauchi, K. Hirota, H. Nakayama, N. Takahashi, T. Isobe, Landscape of the complete RNA chemical modifications in the human 80S ribosome. Nucleic Acids Res. 46, 9289–9298 (2018).

19. S. K. Natchiar, A. G. Myasnikov, H. Kratzat, I. Hazemann, B. P. Klaholz, Visualization of chemical modifications in the human 80S ribosome structure. Nature. 551, 472–477 (2017).

20. G. Keith, G. Dirheimer, The primary structure of rabbit, calf and bovine liver tRNAPhe. BBA Sect. Nucleic Acids Protein Synth. 517, 133–149 (1978).

21. J. A. Kelly, A. N. Olson, K. Neupane, S. Munshi, J. S. Emeterio, L. Pollack, M. T. Woodside, J. D. Dinman, Structural and functional conservation of the programmed -1 ribosomal frameshift signal of SARS coronavirus 2 (SARS-CoV-2) (2020), doi:10.1074/jbc.AC120.013449.

22. J. Rabl, M. Leibundgut, S. F. Ataide, A. Haag, N. Ban, Crystal structure of the eukaryotic 40S ribosomal subunit in complex with initiation factor 1. Science. 331, 730–736 (2011).

23. S. Takyar, R. P. Hickerson, H. F. Noller, mRNA helicase activity of the ribosome. Cell. 120, 49–58 (2005).

24. Z. Lin, R. J. C. Gilbert, I. Brierley, Spacer-length dependence of programmed-1 or-2 ribosomal frameshifting on a U6A heptamer supports a role for messenger RNA (mRNA) tension in frameshifting. Nucleic Acids Res. 40, 8674–8689 (2012).

25. H. Amiri, H. F. Noller, Structural evidence for product stabilization by the ribosomal mRNA helicase. RNA. 25, 364–375 (2019).

26. N. Caliskan, V. I. Katunin, R. Belardinelli, F. Peske, M. V. Rodnina, Programmed -1 frameshifting by kinetic partitioning during impeded translocation. Cell. 157, 1619–1631 (2014).

27. S. J. Park, Y. G. Kim, H. J. Park, Identification of rna pseudoknot-binding ligand that inhibits the - 1 ribosomal frameshifting of SARS-coronavirus by structure-based virtual screening. J. Am. Chem. Soc. 133, 10094–10100 (2011).

28. K. Neupane, S. Munshi, M. Zhao, D. B. Ritchie, S. M. Ileperuma, M. T. Woodside, Anti-Frameshifting Ligand Active against SARS Coronavirus-2 Is Resistant to Natural Mutations of the Frameshift-Stimulatory Pseudoknot. J. Mol. Biol. (2020), doi:10.1016/j.jmb.2020.09.006.

29. M. Bouvet, A. Lugari, C. C. Posthuma, J. C. Zevenhoven, S. Bernard, S. Betzi, I. Imbert, B. Canard, J. C. Guillemot, P. Lécine, S. Pfefferle, C. Drosten, E. J. Snijder, E. Decroly, X. Morelli, Coronavirus Nsp10, a critical co-factor for activation of multiple replicative enzymes. J. Biol. Chem. 289, 25783–25796 (2014).

30. E. C. Smith, J. B. Case, H. Blanc, O. Isakov, N. Shomron, M. Vignuzzi, M. R. Denison, Mutations in Coronavirus Nonstructural Protein 10 Decrease Virus Replication Fidelity. J. Virol. 89, 6418–6426 (2015).

31. J. S. Joseph, K. S. Saikatendu, V. Subramanian, B. W. Neuman, A. Brooun, M. Griffith, K. Moy, M. K. Yadav, J. Velasquez, M. J. Buchmeier, R. C. Stevens, P. Kuhn, Crystal Structure of Nonstructural Protein 10 from the Severe Acute Respiratory Syndrome Coronavirus Reveals a Novel Fold with Two Zinc-Binding Motifs. J. Virol. 80, 7894–7901 (2006).

32. H. R. Harrington, M. H. Zimmer, L. M. Chamness, V. Nash, W. D. Penn, T. F. Miller, S. Mukhopadhyay, J. P. Schlebach, Cotranslational folding stimulates programmed ribosomal frameshifting in the alphavirus structural polyprotein. J. Biol. Chem. 295 (2020), 6798–6808.

33. H. Nakatogawa, K. Ito, The ribosomal exit tunnel functions as a discriminating gate. Cell. 108, 629–636 (2002).

34. D. H. Goldman, C. M. Kaiser, A. Milin, M. Righini, I. Tinoco, C. Bustamante, Mechanical force releases nascent chain-mediated ribosome arrest in vitro and in vivo. Science. 348, 457–460 (2015).

35. G. Kramer, D. Boehringer, N. Ban, B. Bukau, The ribosome as a platform for co-translational processing, folding and targeting of newly synthesized proteins. Nat. Struct. Mol. Biol. 16 (2009), pp. 589–597.

36. K. Döring, N. Ahmed, T. Riemer, H. G. Suresh, Y. Vainshtein, M. Habich, J. Riemer, M. P. Mayer, E. P. O’Brien, G. Kramer, B. Bukau, Profiling Ssb-Nascent Chain Interactions Reveals Principles of Hsp70-Assisted Folding. Cell. 170, 298–311.e20 (2017).

37. M. Lazarczyk, M. Favre, Role of Zn2+ Ions in Host-Virus Interactions. J. Virol. 82, 11486–11494 (2008).

38. A. Sharma, M. Mariappan, S. Appathurai, R. S. Hegde, In vitro dissection of protein translocation into the mammalian endoplasmic reticulum. Methods Mol. Biol. 619, 339–363 (2010).

39. S. Q. Zheng, E. Palovcak, J. P. Armache, K. A. Verba, Y. Cheng, D. A. Agard, MotionCor2: Anisotropic correction of beam-induced motion for improved cryo-electron microscopy. Nat. Methods. 14 (2017), pp. 331–332.

40. K. Zhang, Gctf: Real-time CTF determination and correction. J. Struct. Biol. 193, 1–12 (2016).

41. J. Zivanov, T. Nakane, B. O. Forsberg, D. Kimanius, W. J. H. Hagen, E. Lindahl, S. H. W. Scheres, New tools for automated high-resolution cryo-EM structure determination in RELION-3. Elife. 7 (2018), doi:10.7554/eLife.42166.

42. A. Punjani, J. L. Rubinstein, D. J. Fleet, M. A. Brubaker, CryoSPARC: Algorithms for rapid unsupervised cryo-EM structure determination. Nat. Methods. 14, 290–296 (2017).

43. A. Punjani, D. Fleet, 3D Variability Analysis: Directly resolving continuous flexibility and discrete heterogeneity from single particle cryo-EM images, 1–25 (2020).

44. S. Shao, J. Murray, A. Brown, J. Taunton, V. Ramakrishnan, R. S. Hegde, Decoding Mammalian Ribosome-mRNA States by Translational GTPase Complexes. Cell. 167, 1229–1240.e15 (2016).

45. E. F. Pettersen, T. D. Goddard, C. C. Huang, G. S. Couch, D. M. Greenblatt, E. C. Meng, T. E. Ferrin, UCSF Chimera - A visualization system for exploratory research and analysis. J. Comput. Chem. 25, 1605–1612 (2004).

46. P. Emsley, B. Lohkamp, W. G. Scott, K. Cowtan, Features and development of Coot. Acta Crystallogr. Sect. D Biol. Crystallogr. 66, 486–501 (2010).

47. D. Liebschner, P. V. Afonine, M. L. Baker, G. Bunkoczi, V. B. Chen, T. I. Croll, B. Hintze, L. W. Hung, S. Jain, A. J. McCoy, N. W. Moriarty, R. D. Oeffner, B. K. Poon, M. G. Prisant, R. J. Read, J. S. Richardson, D. C. Richardson, M. D. Sammito, O. V. Sobolev, D. H. Stockwell, T. C. Terwilliger, A. G. Urzhumtsev, L. L. Videau, C. J. Williams, P. D. Adams, Macromolecular structure determination using X-rays, neutrons and electrons: Recent developments in Phenix. Acta Crystallogr. Sect. D Struct. Biol. 75, 861–877 (2019).

48. A. W. Schüttelkopf, D. M. F. Van Aalten, PRODRG: A tool for high-throughput crystallography of protein-ligand complexes. Acta Crystallogr. Sect. D Biol. Crystallogr. 60, 1355–1363 (2004).

49. V. B. Chen, W. B. Arendall, J. J. Headd, D. A. Keedy, R. M. Immormino, G. J. Kapral, L. W. Murray, J. S. Richardson, D. C. Richardson, MolProbity: All-atom structure validation for macromolecular crystallography. Acta Crystallogr. Sect. D Biol. Crystallogr. 66, 12–21 (2010).

50. F. Sievers, A. Wilm, D. Dineen, T. J. Gibson, K. Karplus, W. Li, R. Lopez, H. McWilliam, M. Remmert, J. Söding, J. D. Thompson, D. G. Higgins, Fast, scalable generation of high-quality protein multiple sequence alignments using Clustal Omega. Mol. Syst. Biol. 7 (2011), doi:10.1038/msb.2011.75.

51. X. Robert, P. Gouet, Deciphering key features in protein structures with the new ENDscript server. Nucleic Acids Res. 42 (2014), doi:10.1093/nar/gku316.

52. G. Loughran, M. T. Howard, A. E. Firth, J. F. Atkins, Avoidance of reporter assay distortions from fused dual reporters. RNA. 23, 1285–1289 (2017).

53. B. W. Dyer, F. A. Ferrer, D. K. Klinedinst, R. Rodriguez, A noncommercial dual luciferase enzyme assay system for reporter gene analysis. Anal. Biochem. 282, 158–161 (2000).

54. Virology Methods Manual (Elsevier, 1996).

55. J. Schindelin, I. Arganda-Carreras, E. Frise, V. Kaynig, M. Longair, T. Pietzsch, S. Preibisch, C. Rueden, S. Saalfeld, B. Schmid, J. Y. Tinevez, D. J. White, V. Hartenstein, K. Eliceiri, P. Tomancak, A. Cardona, Fiji: An open-source platform for biological-image analysis. Nat. Methods. 9 (2012), pp. 676–682.

56. J. Mutterer, E. Zinck, Quick-and-clean article figures with FigureJ. J. Microsc. 252, 89–91 (2013).

57. T. D. Goddard, C. C. Huang, E. C. Meng, E. F. Pettersen, G. S. Couch, J. H. Morris, T. E. Ferrin, UCSF ChimeraX: Meeting modern challenges in visualization and analysis. Protein Sci. 27, 14–25 (2018).

58. G. Cardone, J. B. Heymann, A. C. Steven, One number does not fit all: Mapping local variations in resolution in cryo-EM reconstructions. J. Struct. Biol. 184, 226–236 (2013).

59. V. Chandrasekaran, S. Juszkiewicz, J. Choi, J. D. Puglisi, A. Brown, S. Shao, V. Ramakrishnan, R. S. Hegde, Mechanism of ribosome stalling during translation of a poly(A) tail. Nat. Struct. Mol. Biol. 26, 1132–1140 (2019).

