## Supplementary material for "Structural basis of ribosomal frameshifting during translation of the SARS-CoV-2 RNA genome": Materials and Methods and all supplementary material

**This PDF file includes:**

Materials and Methods
Figs. S1 to S5
Tables S1 to S2

### Materials and Methods

Generation of DNA templates and *in vitro* transcription of the mRNA

Constructs used for testing and generating ribosomal complexes *in vitro* contain the following elements: A linear DNA template containing a T7 promoter and Kozak sequence upstream of a sequence encoding an N-terminal 3x FLAG-tag followed by a 290 nt linker, Wild type (WT) or mutant SARS-CoV-2 frameshift context and the majority of *nsp12* was synthesized as a gene fragment by Genscript. The SARS-CoV-2 WT/mutant frameshift context sequences in turn consist of the following elements in sequential order: The final 115 nucleotides of sequence encoding the C-terminal end of Nsp10; the coding sequence of Nsp11/Nsp12 (which includes the so-called 'attenuator loop') up to the Phe 'UUU' codon that just precedes the frameshift site; WT ('UUA AAC') or mutant ('UUA UAA') frameshift sites; the WT frameshift stimulatory pseudoknot sequence; 540 nucleotides of sequence encoding Nsp12 after the frameshift site.

The entire fragment was PCR amplified from opposite ends with complementary primers using Q5 DNA Polymerase (NEB) and purified using the QIAquick PCR Purification kit (Qiagen). *In vitro* transcription was performed using the Ampliscribe T7 Flash system (Lucigen), followed by lithium chloride precipitation to purify the synthesized RNA.

Sequence of synthesized DNA (Gene Universal, USA) used to test WT SARS-CoV-2 frameshifting efficiency in RRL (Frameshift context underlined):

GATATTCTAAGTGACGTTACGCTAGTGATCGCCGTAATACGACTCACTATAGGCAACAACAA
CAAACATTTGCTTCTGACACAACTGTGTTCACTAGCAACCTCAAACAGACACCATGGACTAC
AAAGACCACGACGGTGATTATAAAGATCACGACATCGATTACAAGGACGACGACGACAAGT
CCAAGGAGCCGCTTCGGCCACGGTGCCGCCCCATCAACGCCACCCTGGCTGTGGAGAAGGA
GGGCTGCCCCGTGTGCATCACCGTCAACACCACCATCTGTGCCGGCTACTGCCCCACCGCAA
CCCGCGTGCTGCAGGGGGTCCTGCCGGCCCTGCCTCAGGTGGTGTGCAACTACCGCCGGTCC
GTAACCCACCGTATTCTTACCGTTCCGATTGCCCAAGATCAAGTGGGCGCATACTATCAGCA
ACCAGGTCAGCAGAACGCCACCTGGATTGTGCCACCAGGGCCAACTTGTGCTAATGACCCTG
TGGGTTTTACACTTAAAAACACAGTCTGTACCGTCTGCGGTATGTGGAAAGGTTATGGCTGT
AGTTGTGATCAACTCCGCGAACCCATGCTTCAGTCAGCTGATGCACAATCGTTTTTAAACGG
GTTTGCGGTGTAAGTGCAGCCCGTCTTACACCGTGCGGCACAGGCACTAGTACTGATGTCGT
ATACAGGGCTTTTGACATCTACAATGATAAAGTAGCTGGTTTTGCTAAATTCCTAAAAACTA
ATTGTTGTCGCTTCCAAGAAAAGGACGAAGATGACAATTTAATTGATTCTTACTTTGTAGTTA
AGAGACACACTTTCTCTAACTACCAACATGAAGAAACAATTTATAATTTACTTAAGGATTGT
CCAGCTGTTGCTAAACATGACTTCTTTAAGTTTAGAATAGACGGTGACATGGTACCACATAT
ATCACGTCAACGTCTTACTAAATACACAATGGCAGACCTCGTCTATGCTTTAAGGCATTTTG
ATGAAGGTAATTGTGACACATTAAAAGAAATACTTGTCACATACAATTGTTGTGATGATGAT

TATTTCAATAAAAAGGACTGGTATGATTTTGTAGAAAACCCAGATATATTACGCGTATACGCCAACTTAGGTGAACGTGTACGCCAAGCTTTG

Sequence of synthesized DNA (Gene Universal, USA) used to test 0 frame stop codon mutation of SARS-CoV-2 frameshifting efficiency in RRL (mutant frameshift context underlined):
GATATTCTAAGTGACGTTACGCTAGTGATCGCCGTAATACGACTCACTATAGGCAACAACAACAAACATTTGCTTCTGACACAACTGTGTTCACTAGCAACCTCAAACAGACACCATGGACTACAAAGACCACGACGGTGATTATAAAGATCACGACATCGATTACAAGGACGACGACGACAAGTCCAAGGAGCCGCTTCGGCCACGGTGCCGCCCCATCAACGCCACCCTGGCTGTGGAGAAGGAGGGCTGCCCCGTGTGCATCACCGTCAACACCACCATCTGTGCCGGCTACTGCCCCACCGCAACCCGCGTGCTGCAGGGGGTCCTGCCGGCCCTGCCTCAGGTGGTGTGCAACTACCGCCGGTCCGTAACCCACCGTATTCTTACCGTTCCGATTGCCCAAGATCAAGTGGGCGCATACTATCAGCAACCAGGTCAGCAGAACGCCACCTGGATTGTGCCACCAGGGCCAACTTGTGCTAATGACCCTGTGGGTTTTACACTTAAAAACACAGTCTGTACCGTCTGCGGTATGTGGAAAGGTTATGGCTGTAGTTGTGATCAACTCCGCGAACCCATGCTTCAGTCAGCTGATGCACAATCGTTTTTATAAGGGTTTGCGGTGTAAGTGCAGCCCGTCTTACACCGTGCGGCACAGGCACTAGTACTGATGTCGTATACAGGGCTTTTGACATCTACAATGATAAAGTAGCTGGTTTTGCTAAATTCCTAAAAACTAATTGTTGTCGCTTCCAAGAAAAGGACGAAGATGACAATTTAATTGATTCTTACTTTGTAGTTAAGAGACACACTTTCTCTAACTACCAACATGAAGAAACAATTTATAATTTACTTAAGGATTGTCCAGCTGTTGCTAAACATGACTTCTTTAAGTTTAGAATAGACGGTGACATGGTACCACATATATCACGTCAACGTCTTACTAAATACACAATGGCAGACCTCGTCTATGCTTTAAGGCATTTTGATGAAGGTAATTGTGACACATTAAAAGAAATACTTGTCACATACAATTGTTGTGATGATGATTATTTCAATAAAAAGGACTGGTATGATTTTGTAGAAAACCCAGATATATTACGCGTATACGCCAACTTAGGTGAACGTGTACGCCAAGCTTTGTAATAGTCATAGAGGAT

Purification of eRF1(AAQ)

Mutant eRF1(AAQ) placed in a vector containing an N-terminal 6x His-tag followed by a GST-tag and TEV protease cleavage site was expressed from a pET-24 d(+) vector. The plasmid was transformed into *E. coli* BL21 (DE3) cells under Kanamycin selection, and cells were grown in 2× YT medium at 37 °C. At an optical density at 600 nm (OD600) of 0.8, cultures were shifted to 18 °C and induced with IPTG added to a final concentration of 1 mM. After 16 h, cells were collected by centrifugation, resuspended in lysis buffer (50 mM $NaH_2PO_4$, 300 mM NaCl, 10 mM imidazole, 5 mM β-mercaptoethanol, pH 8.0) and lysed using a cell disrupter (Constant Systems). The lysate was cleared by centrifugation for 30 min at 48’000 x g and loaded onto a batch-binding column of Ni-NTA beads (Agarose Bead Technologies). After binding at 4 °C on a nutator, beads were washed with 10 column volumes of lysis buffer containing 45 mM imidazole, and proteins were eluted in 1.5 column volumes of lysis buffer containing 300 mM imidazole. Eluted proteins were dialyzed overnight in the presence of TEV protease (produced in-house) and passed over Ni-NTA beads. Tag-free eRF1(AAQ) was collected in the flowthrough, and loaded on to a Superdex 75 16/60 column in storage buffer (50 mM HEPES, pH 7.4, 150 mM KOAc, 5 mM $Mg(OAc)_2$, 10 mM imidazole, 10% glycerol, 1 mM DTT).

<u>*In vitro* translation reaction and RNC purification</u>

Translationally active RRL was generated from untreated rabbit reticulocyte lysate (Green Hectares, USA) as described in (*38*) with certain modifications. The final translation reaction (per 20 µL) contained 11.4 µL activated nuclease-treated and activated RRL, 200 ng/µL mRNA, 1.2 mM $MgCl_2$, 0.5 mM DTT, 0.4 mM spermidine and eRF1(AAQ) at a final concentration of 3.5 µM to trap translating ribosomes. Reactions were incubated for 25 min at 32 °C. 4 mL translation reactions were chilled on ice for 10 mins to halt translation, and HEPES KOH pH 7.5 was added to a final concentration of 50 mM. Chilled translation reactions were directly incubated with 400 µL of packed anti-FLAG M2 beads (Sigma) for 2 h at 4 °C with gentle mixing. The beads were then washed with 4 ml of 50 mM HEPES, pH 7.4, 100 mM KOAc, 5 mM $Mg(OAc)_2$, 0.1% Triton X-100, 1 mM DTT; 4 ml of 50 mM HEPES, pH 7.4, 250 mM KOAc, 5mM $Mg(OAc)_2$, 0.5% Triton X-100, 1 mM DTT and 6 ml of RNC buffer (50mM HEPES, pH 7.4, 100 mM KOAc, 5 mM $Mg(OAc)_2$, 1mM DTT). RNCs were eluted after 4 sequential 10 min incubations at room temperature in RNC buffer that contained 0.2 mg/mL 3x FLAG peptide (Sigma). The elutions were combined and centrifuged at 186,000 x g at 4 °C for 2 h in a TLA 55 rotor (Beckman Coulter). Supernatant was discarded and the pellet was resuspended in RNC buffer at a concentration of 80 nM. At each step of translation reactions and purification, aliquots were taken to perform Western blots using an anti-FLAG antibody (Sigma Cat. No. A8592).

<u>Cryo-electron microscopy, sample preparation and data collection</u>

To prepare cryo-EM grids using a Vitrobot (ThermoFisher Scientific), 5 µL of sample were applied to Quantifoil R2/2 holey carbon copper grids (Quantifoil Micro Tool), which were covered with a sheet of continuous carbon and glow-discharged for 15 s at 15 mA using a easiGlow Discharge cleaning system (Pelco) beforehand. The sample was incubated for 30 s in the Vitrobot chamber, which was kept at 4 °C and 100% humidity. The excess sample was then blotted for 6 to 10 s and immediately plunged into a mixture of ethane:propane (1:2). Grids were loaded into a Titan Krios cryo-transmision electron microscope (ThermoFisher Scientific) operating at 300 kV, and data collection was performed using a GIF Quantum LS energy filter (Gatan)-mounted K3 direct electron camera (Gatan) in counting and super-resolution mode. The microscope was used with a 81'000 x nominal magnification, resulting in a physical pixel size of 1.08 Å/pixel and therefore a super-resolution pixel of 0.54 Å/pixel. Both data collections were set up with the EPU program (Thermo Fisher Scientific), taking advantage of the Aberration-Free Image Shift (AFIS) mode. 40-frame movies were recorded with a total dose of 60 $e^{-}$ * $Å^{-2}$, and the defocus was set to change between -0.6 and -3 µm with 0.3 µm increments. The energy filter slit width was set to 20 eV during exposure.

Cryo-electron microscopy data processing (fig. S1)

Approximately 10’000 movies were processed with MotionCor2 (*39*) to dose-weight the radiation damage, apply the gain correction, correct for motion during the exposure, and bin twice the super-resolution micrographs. GCTF (*40*) was then used to estimate the Contrast Transfer Functions (CTF) of the motion-corrected and dose-weighted micrographs.

Based on both the quality of the micrographs and their respective CTF, 9’580 micrographs were selected for further processing. 1’897’486 particles were picked using the Laplacian-of-Gaussian-based method in Relion3.1(*41*). Those particles were extracted using Relion3.1 and then imported into cryoSPARC v2 (*42*). After 2D classification using cryoSPARC v2 to select good particles, they were first refined into a 3D structure, followed by a two-step classification using cryoSPARC v2 heterogeneous refinement and a variability analysis (*43*). Out of those classifications, 695’501 particles containing a visible pseudoknot were selected for final refinement, resulting in a reconstruction with an overall resolution of 2.2 Å (fig. S1).

Those particles were then further classified in Relion3.1 using a mask around the pseudoknot, which resulted in a class containing 171’706 particles with the three stems of the pseudoknot well visible. Those particles were refined in cryoSPARC v2 to reach an overall resolution of 2.4 Å (fig. S1).

General model building

As a starting model, the small subunit head, body and the large subunit of the 3.3Å structure of a rabbit ribosomal elongation complex (PDB 5LZS (*44*)) were rigid body docked into the 2.2 Å cryo-EM map using CHIMERA (*45*). While the ribosomal proteins could be readily readjusted in COOT (*46*), obvious discrepancies in the rRNA sequences lead us to retrieve updated full-length sequences for the 28S and 5.8S rRNAs, the 18S rRNA and the 5S rRNA from the *Oryctolagus cuniculus* Transcriptome Shotgun Assembly (TSA) at the GeneBank (entries GBCN01009604.1, GBCT01000564.1 and GBCM01014045.1, respectively). The model was revised and completed using a sharpened 2.2 Å cryo-EM map in combination with the further classified 2.4Å cryo-EM map. While the further classified map revealed the features of the COVID-19 mRNA with the pseudoknot, the nascent chain, and tails of peripheral proteins and rRNA expansion segments, the higher resolved map showed clear additional densities for the rRNA and tRNA modifications, bound spermine/spermidine ligands, ions such as hexa-coordinated $Mg^{2+}$, and the ordered solvent (fig. S3). The strictly octahedrally coordinated magnesium ions were built according to their star-shaped appearance, while the nature of numerous other ions remained unassigned due to incomplete or asymmetric coordination shells. Automated water picking was performed using the phenix.douse tool recently developed for ordered solvent building into high-resolution cryo-EM maps and implemented in PHENIX (*47*), followed by manual examination.

Building of the modified rRNA and tRNA(Phe) residues

In an attempt to make use of the modified rRNA residues described in a cryo-EM structure of the human ribosome (PDB 6EK0 (*19*)), we superimposed the human rRNA onto our structure. However, it turned out that the modification pattern disagreed with our high-resolution EM-map to a considerable extent. In contrast, when we used for model building the full set of human rRNA modifications recently established by quantitative mass spectroscopy, we observed that the modification pattern coincided with our maps remarkably well, supporting the existence of eukaryotic-typical rather than human-specific rRNA modifications (*18*). Although the difference between pseudouridines and uridines cannot be directly established from the overall shape of the base, we saw a water molecule bound to the N1 atom of the pseudouridine ring in many positions, while in uridine such a hydrogen bond does not exist. Therefore, apart from all modifications comprising the easily distinguishable additional groups, we decided to also include the complete set of pseudouridines in our model, which may serve as a template for other mammalian high-resolution structures in the future. For modelling of the P-site tRNA(Phe), we followed the sequence and modification pattern of mammalian tRNA(Phe) determined by Keith and Dirheimer (*20*), which could be built without any obvious discrepancy into our maps. As the E-site tRNA was less-well resolved and its identity could not be established from the maps, we modeled a generic tRNA and left the residues unassigned.

Real space refinement

The completed structure was refined for five cycles into the higher resolution (including ordered solvent) or further classified (excluding waters not involved in $Mg^{2+}$ hexa-coordination) maps, respectively, using real space refinement in PHENIX version 1.18 (*47*). Custom restraints for modified rRNA residues were generated using PRODRG (*48*) or exported from COOT (*46*), and protein secondary structure and Ramachandran as well as RNA base pair and stacking restraints were applied throughout to maintain good model geometry also in less-well-ordered areas at the periphery of the 80S maps. Remaining discrepancies between the model and the maps were detected and corrected using real space difference maps, followed by two additional cycles of refinement as described above. The structures were validated using MOLPROBITY (*49*). The resulting models display excellent geometries and map correlations (table S1), and the resolution of the model *vs.* map FSCs at a value of 0.5 coincide well with the resolution determined between the map half-sets at the FSC=0.143 criterion (fig. S2B).

Alignment of SARS2 with sequences of related coronaviruses

The sequences for the alignments shown in Fig. 4G were obtained from the UniProt database (https://www.uniprot.org/) and correspond to the polyprotein translated from ORF1a before the frameshift event takes place. The following sequences were used: P0DTC1 (SARS2), P0C6T7 (Bat coronavirus Rp3), P0C6U8 (SARS), P0C6F5 (Bat coronavirus 279/2005), A0A0K1Z0N1 (Bat SARS-like coronavirus YNLF_31C), R9QTH2 (Bat coronavirus Cp/Yunnan2011), P0C6T6 (Bat coronavirus HKU9), K9N638 (MERS), U5LR11 (*Betacoronavirus Erinaceus*/VMC/DEU/2012). The sequences were aligned with ClustalOmega (https://www.ebi.ac.uk/Tools/msa/clustalo/) (*50*) and visualized with ESPript (http://espript.ibcp.fr) (*51*).

Structure-guided mutagenesis experiments

Dual luciferase expression constructs were generated by either 1-step or 2-step PCR using primer sequences outlined in table S2, during which 5' *Xho*I and 3' *Bam*HI restriction sites were incorporated. PCR amplicons were digested with *Xho*I / *Bam*HI and cloned into a *Psp*XI / *Bgl*II -digested pSGDlucV3.0 vector (Addgene 119760, (*52*)).

HEK293T cells (ATCC) were maintained in DMEM supplemented with 10% FBS, 1 mM L-glutamine and antibiotics. HEK293T cells were transfected with Lipofectamine 2000 reagent (Invitrogen) using the 1-day protocol, in which suspended cells are added directly to the DNA complexes in half-area 96-well plates. To each well, 25 ng of each plasmid plus 0.2 µl Lipofectamine 2000 in 25 µl Opti-Mem (Gibco) were added. The transfecting DNA complexes in each well were incubated with $4 \times 10^4$ cells suspended in 50 µl DMEM + 10% FBS at 37 °C in 5% $CO_2$ for 24 h.

For the *in vitro* translation reactions, Plasmid DNAs (50 ng) were used as templates in 5 µl reactions of the RRL TNT® T7 Quick Coupled Transcription/Translation system (Promega) supplemented with 1 mM methionine (25 µM final concentration). Reactions were incubated at 30 °C for 90 min.

For the Dual Luciferase Assay, relative light units were measured on a Veritas Microplate Luminometer with two injectors (Turner Biosystems). Transfected cells were lysed in 15 µl of 1 × passive lysis buffer (PLB), and light emission was measured following injection of 50 µl of either Renilla or firefly luciferase substrate (*53*). For *in vitro* translation reactions, 45 µl of 1 × PLB was added to each 5 µl reaction, and luciferase activities were assayed from 10 µl.

Frameshifting efficiencies (% frameshifting) were determined by calculating relative luciferase activities (firefly/Renilla) from test constructs and dividing by relative luciferase activities from replicate wells of matched in-frame control constructs. The number of biological replicates for each experiment are indicated in each figure legend. For all Boxplots center line medians are shown; box limits indicate the 25$^{th}$ and 75$^{th}$ percentiles as determined by the R software; whiskers extend 1.5 times the interquartile range from the 25$^{th}$ and 75$^{th}$ percentiles, outliers are represented by dots.

Synthesis of ethyl 2-(4-((2-methylthiazol-4-yl)methyl)-1,4-diazocane-1-carboxamido)benzoate (MTDB)

The RNA pseudoknot-binding ligand ethyl 2-(4-((2-methylthiazol-4-yl)methyl)-1,4-diazocane-1-carboxamido)benzoate was prepared from commercially obtained materials using the following procedure. Under a nitrogen atmosphere, 4-((1,4-diazocan-1-yl)methyl)-2-methylthiazole (110 mg, 0.51 mmol) was charged to a 10 mL oven dried round bottom flask equipped with magnetic stirring. Anhydrous DMF (2.0 mL) was added and the solution cooled to 0 °C using an ice bath. Ethyl 2-isocyanatobenzoate (130 mg, 0.70 mmol) was dissolved in anhydrous DMF (2.0 mL) and added to the reaction dropwise. The reaction was stirred at 0 °C for 1 h then diluted with water (40 mL). The product was extracted in ethyl acetate (4 x 15 mL). The combined organic phases were washed with water (4 x 20 mL) and brine (2 x 20 mL) then dried over an excess of anhydrous magnesium sulfate. The resulting solution was concentrated under reduced pressure to afford the crude product which was purified by flash silica chromatography (100% ethyl acetate) to afford the title compound as a light-yellow oil (170 mg, 0.43 mmol, 84% yield). The product was characterized by NMR spectroscopy (1H, 13C, COSY, HSQC, HMBC on a Bruker AV-400 and -500 MHz spectrometer), IR spectroscopy (thin film, Jasco FT/IR-4100) and HRMS (measured by the mass spectrometry service of the ETH Zürich Laboratorium für Organische Chemie on a Bruker Daltonics maXis ESI-QTOR spectrometer). The ligand was stored in a sealed vial at 4 °C.

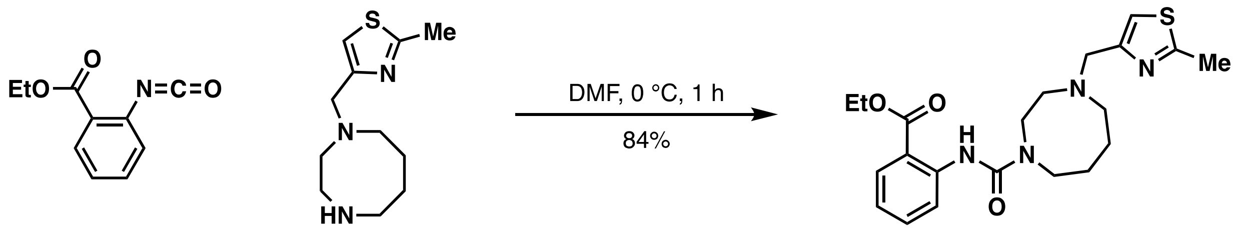


Testing the effect of MTDB on SARS-CoV-2 infected cells

VeroE6 cells (kindly provided by Doreen Muth, Marcel Müller and Christian Drosten, Charité, Berlin, Germany) were propagated in Dulbecco's modified EMEM (DMEM), supplemented with 10% heat inactivated fetal bovine serum, 1% nonessential amino acids, 100 µg/mL of streptomycin and 100 IU/mL of penicillin, and 15 mM of HEPES. Cells were maintained at 37 °C in a humidified incubator with 5% $CO_2$. Cells were infected using SARS-CoV-2 (SARS-CoV-2/München-1.1/2020/929, kindly provided by Daniela Niemeyer, Marcel Müller and Christian Drosten, Charité, Berlin, Germany).

Antiviral analysis

Cells were plated to 20,000 cells per 96 well 24 h prior to infection. Cells were infected with SARS-CoV-2 at a multiplicity of infection (MOI) of 0.01 for 1.5 h at 37 °C and washed 3 times with PBS. MTDB (or respective volumes of its 20 % DMSO/100 mM HCl solvent) was added to cells in following concentrations: 0 µM, 5 µM, 10 µM, 20 µM, 50 µM, 100 µM, 150 µM and 200 µM. 24 hours post infection virus-containing supernatant was serially diluted, and the 50% tissue culture infectious dose ($TCID_{50}$) per milliliter was displayed using Crystal Violet and calculated by the Spearman-Kärber algorithm after 72 h as described (*54*). Cytotoxic effects of MTDB or its 20 % DMSO/100 mM HCl solvent were monitored using CytoTox 96® Non-Radioactive Cytotoxicity Assay (Promega).

Immunofluorescence analysis

VeroE6 cells were fixated with 4% formalin. Cells were permeabilized in PBS supplemented with 50 mM $NH_4Cl$, 0.1% (w/v) saponin and 2% (w/v) bovine serum albumin. Cells were immunostained with mouse monoclonal antibody against dsRNA (SCICONS, clone J2). Alexa-Fluor 488-labeled donkey-anti mouse IgG (H+L) (JacksonImmuno) was used as secondary antibody. Images were acquired using an EVOS FL Auto 2 Imaging System, using 10x objective, processed using Fiji software packages (*55*) and assembled using the FigureJ plugin (*56*).

Figure generation

All density and structure representations were generated using UCSF ChimeraX (*57*) Local Resolution estimate were performed within cryoSPARC v2 (*42*), which uses an implementation of BlocRes (*58*).

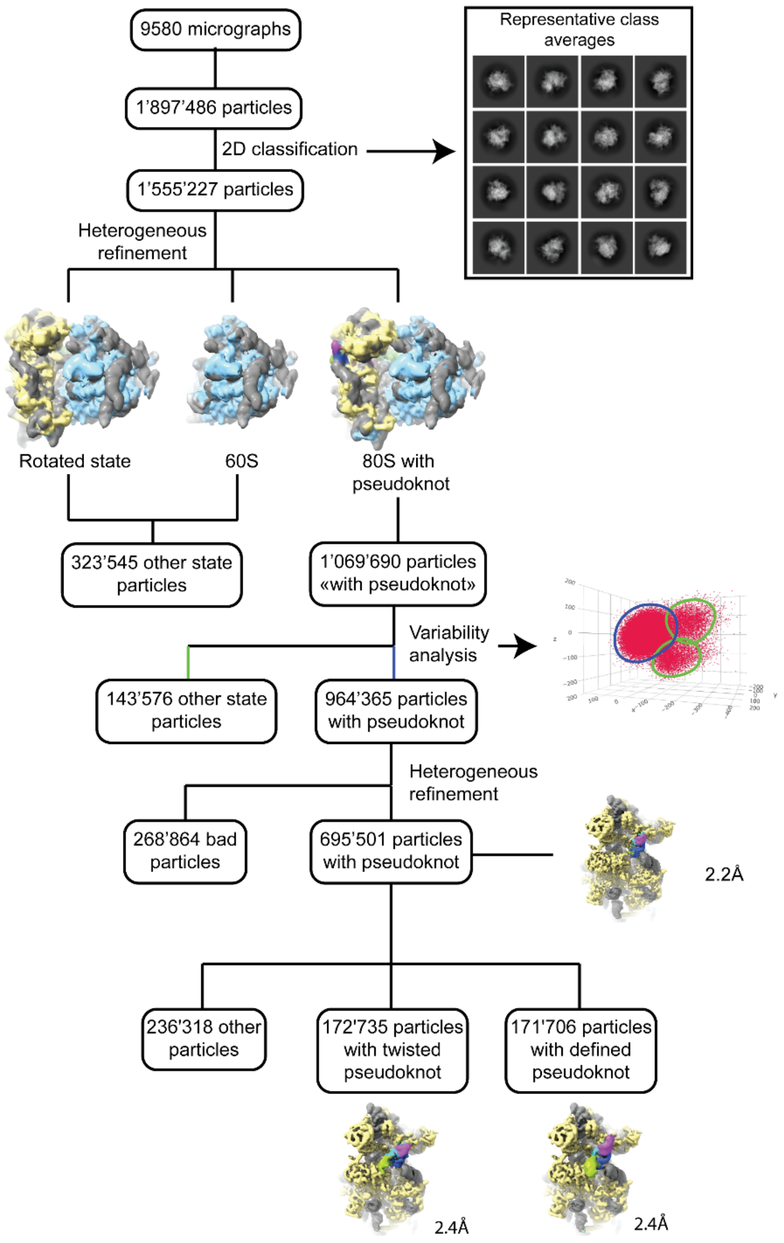


**Fig. S1 Cryo-EM data processing workflow.**

From 9580 micrographs selected for their quality and the quality of their Contrast Transfer Function (CTF), 1’897’486 particles were picked and extracted using Relion3.1 (*41*). The particles were then imported into cryoSPARC v2 (*42*) and processed to remove bad particles, 60S ribosomal subunit and 80S ribosomes in other states. 695’501 selected particles were refined using cryoSPARC v2 to reach an overall resolution of 2.2 Å (fig. S2A-B), corresponding to the Nyquist frequency for our collection pixel size. Further classification using Relion3.1 allowed the separation of a good fractions of particles (171’706) with a clearly defined pseudoknot, which were then refined in cryoSPARC v2 to an overall resolution of 2.4 Å (fig. S2B-C).

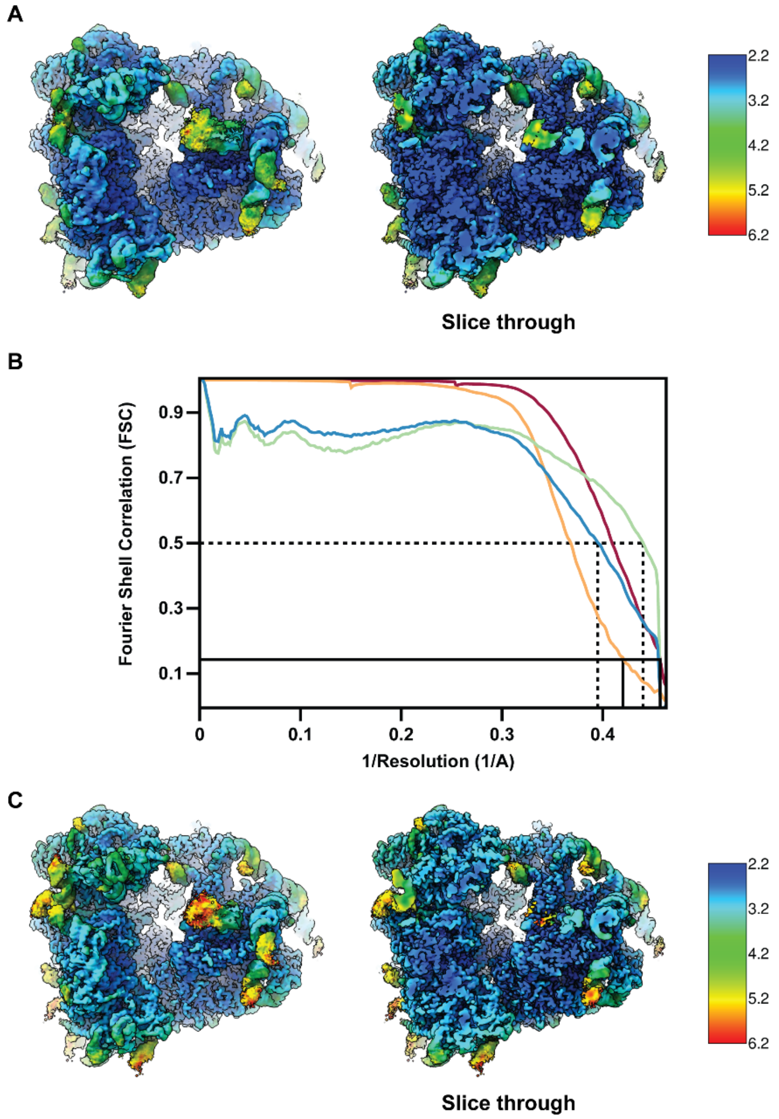


**Fig. S2 Local resolution estimate and cryo-EM FSC.**
(**A**) Local resolution heat map of the high resolution cryo-EM reconstruction with a slice through the density on the right. The local resolution was calculated with the cryoSPARC v2 (*42*) implementation of BlocRes (*58*). The local resolution varies from 2.2Å in the center of the ribosome to roughly 5 to 6 Å for the flexible rRNA expansion and protein segments at the periphery. (**B**) Fourier shell correlations (FSCs) between masked half maps for the high resolution cryo-EM map (red) and the reconstruction of the further classified set of particles (orange). FSCs were also calculated between maps and models for the high resolution (green) and the further classified set (blue). The similar values for the obtained resolutions of map-versus-map and map-versus-model FSCs indicates the absence of overfitting. (**C**). Similar to panel A, but for the reconstruction from further classified set of particles. The resolution varies from roughly 2.5 Å in the center to 5-6 Å for the peripheral rRNA expansion and protein segments and the pseudoknot.

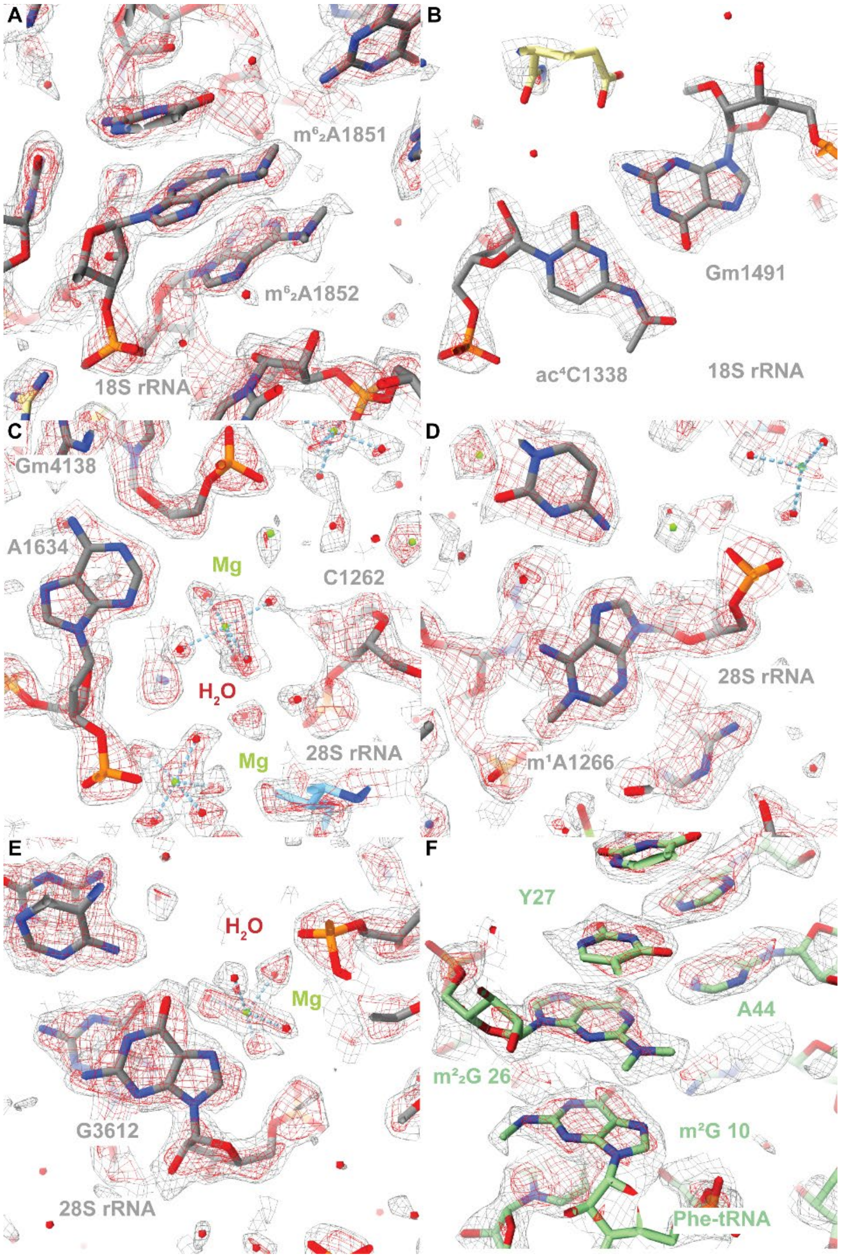


**Fig. S3 Close-up view of rRNA modification and quality of the cryo-EM reconstruction.** The high resolution cryo-EM map is shown at two contour levels as grey and red mesh. (**A**) Close-up view of two $N^6$,$N^6$-dimethyladenosine found at position 1851 and 1852 of the SSU 18S rRNA. (**B**). Close-up view of two modified residues base-pairing in the SSU 18S rRNA: $N^4$-acetylcytidine 1338 and $O^2$-methylguanosine 1491. (**C**) Close-up view of coordinated magnesium ions found in the core of the LSU, for which clear density for the water molecules can be observed. (**D**) Close-up view of the 1-methyladenosine 1266 of the LSU 28S rRNA (**E**) Close-up view of G3612 of the 28S rRNA which was previously reported as 7-propylguanosine 3880 (PDB 6EK0 (*19*)). (**F**) Close-up view of the Phe-tRNA modifications pseudouridine 27, $N^2$,$N^2$-dimethylguanosine 26, and $N^2$-methylguanosine 10.

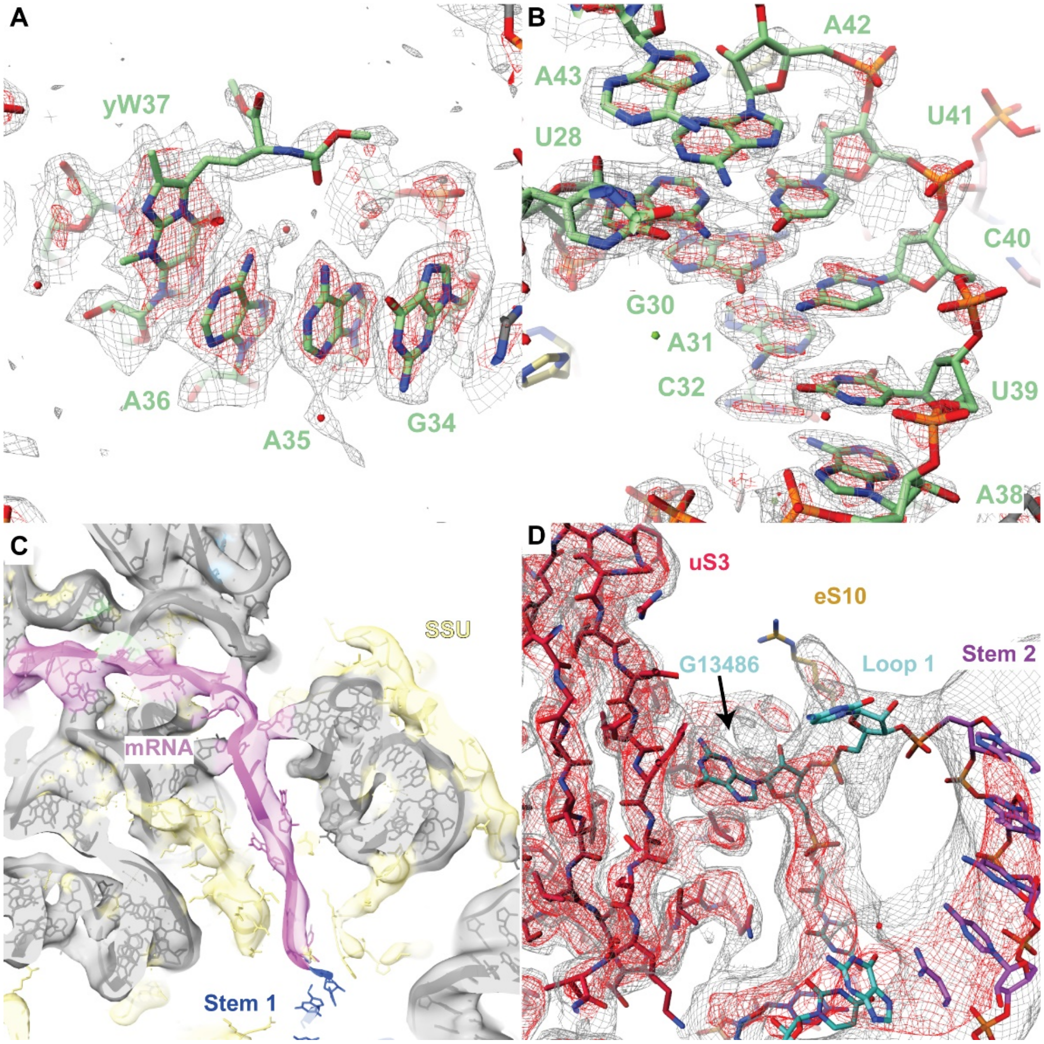


**Fig. S4 Close-up view of features in cryoEM maps.**

Densites shown in panel (A), (B) and (D) are shown at two contour levels as grey and red mesh. (**A**) Close-up view of the anticodon loop of the Phe-tRNA found in the P-site. The hyper-modified wybutosine in position 37 can be clearly identified in the density of the high-resolution cryo-EM reconstruction. (**B**) Close-up view of the anticodon stem-loop shown in the high-resolution cryo-EM map. The tRNA in the P-site could be unambiguously identified as Phe-tRNA based on the purine-pyrimidine pattern of the codon-anticodon (Fig. 1E), its modification pattern (fig. S3), the anticodon stem loop shown here, and the attached amino acid residues of the nascent chain (**C**) Close-up view of the mRNA density seen in the cryo-EM map after further classification. A continuous density can be seen from the P-site codon up to the stem of the pseudoknot, allowing us to set the pseudoknot registry start, which is shifted relative to the previously reported secondary structure diagram (*13*). (**D**) Close-up view of the loop 1 of the pseudoknot, where residue G13486 can be seen being flipped out to contact uS3. The density shown corresponds to the cryo-EM map classified for the pseudoknot.

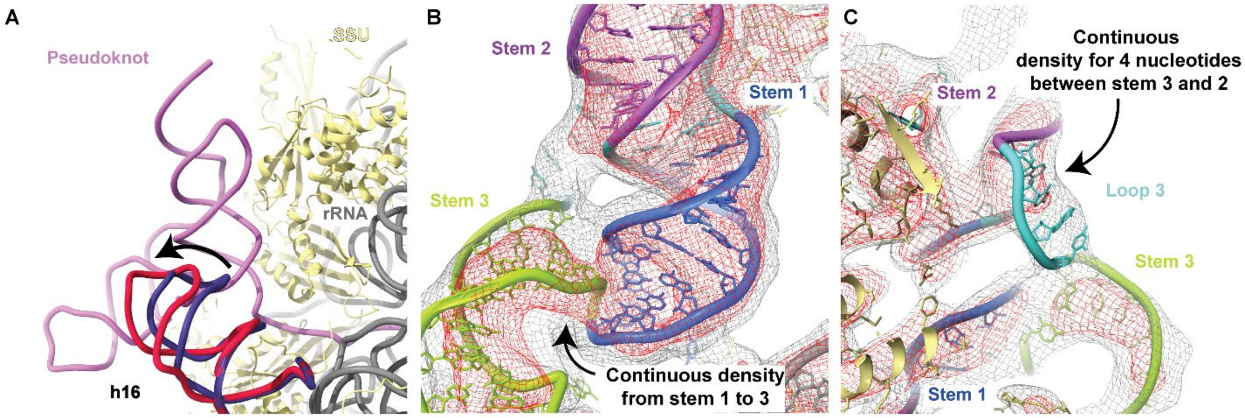


**Fig. S5 Structure of the pseudoknot and interaction with h16.**

(**A**) Close-up view of the pseudoknot (pink) bound to the SSU (yellow proteins and grey rRNA). The pseudoknot can be seen interacting with 18S rRNA helix h16 (red), pushing it outwards compared to translating ribosome (purple, PDB 6SGC (*59*). (**B**) Close-up view of the pseudoknot to highlight the transition between the Stems 1 and 3. The direct connection requires adjustments relative to the previously reported secondary structure diagram and agrees with the altered base-pairing pattern observed for Stem 1 (fig. S4C). (**C**) Close-up view of the other side of the pseudoknot relative to panel B. Loop 3 is well visible and corresponds in length to the newly defined four unpaired nucleotides, in agreement with our new secondary structure diagram.

**Table S1.EM data collection and structure refinement statistics.**

| | 80S – SARS2 pseudoknot (higher resolution map) (EMD-YYYY, PDB XXXX) | 80S – SARS2 pseudoknot (further classified set map) (EMD-WWWW, PDB ZZZZ) |
|---|---|---|
| Data collection and processing | | |
| Magnification | 81’000x (nominal) | |
| Voltage (kV) | 300 | |
| Electron exposure (e–/Å2) | 60 | |
| Defocus range (µm) | 0.6-3 | |
| Pixel size (Å) | 1.08 (super-resolution pixel at 0.54Å/pixel) | |
| Initial particle images (no.) | 1’897’486 | |
| Final particle images (no.) | 695’501 | 171'706 |
| Map resolution at FSC=0.143 (Å) | 2.2 | 2.4 |
| Refinement | | |
| Model resolution at FSC=0.5 (Å) | 2.3 | 2.7 |
| CCmask | 0.81 | 0.82 |
| Map sharpening B factor (Å2) | – 75.8 | – 71.7 |
| Model composition | | |
| Non-hydrogen atoms | 237’931 | |
| Protein residues | 12’093 | |
| RNA residues (modified) | 6050 (220) | |
| Ligands: Zn2+/Mg2+/other ions/SPM/SPD | 8/382/284/4/22 | |
| Waters (strictly coordinated to Mg2+) | 10’155 (1874) | - (1874) |
| B factors min/max/mean (Å2) | | |
| Protein | 1/50/25 | 7/57/27 |
| RNA | 1/78/33 | 13/87/37 |
| Ligand | 3/56/21 | 9/66/23 |
| Water | 1/104/17 | - |
| R.m.s. deviations | | |
| Bond lengths (Å) | 0.002 | 0.002 |
| Bond angles (°) | 0.667 | 0.659 |
| Validation | | |
| MolProbity score | 1.6 | 1.6 |
| Clashscore | 5.88 | 5.77 |
| Poor rotamers (%) | 1.81 | 2.08 |
| Protein | | |
| EM Ringer score | 3.97 | 3.55 |
| Ramachandran plot | | |
| Favored (%) | 96.63 | 97.8 |
| Allowed (%) | 2.33 | 2.15 |
| Disallowed (%) | 0.04 | 0.04 |
| RNA | | |
| Pucker outliers (%) | 0.10 | 0.10 |
| Bond outliers (%) | 0.29 | 0.29 |
| Angle outliers (%) | 0.03 | 0.03 |
| Suite outliers (%) | 14.5 | 14.7 |

**Table S2. List of primers used for mutagenesis experiments**

| Primer name | Primer sequence (5'-3') |
|---|---|
| SARS CoV2 WT S XhoI | ATAACTCGAGACCAACTTGTGCTAATGACCCTGTG |
| SARS CoV2 WT AS BamHI | ATAAGGATCCATTGTAGATGTCAAAAGCCC |
| G-A S | GCGGTGTAAGTACAGCCCGTCTTACAC |
| G-A AS | GTGTAAGACGGGCTGTACTTACACCGC |
| G-C S | GCGGTGTAAGTCCAGCCCGTCTTACAC |
| G-C AS | GTGTAAGACGGGCTGGACTTACACCGC |
| del L1 S | GGGTTTGCGGTGTAAGAGCCCGTCTTACACCG |
| del L1 AS | ACGGTGTAAGACGGGCTCTTACACCGCAAACCC |
| Bulged A deletion AS | ATAAGGATCCGATTGTAGATGTCAAAAGCCCGTATACGACATCAGTAC |
| del L3 AS | TGTAGATGTCAAAAGCCCATACGACATCAGTAC |
| plus 6 stop S | GCGGTGAAAGTGTAGCCCGTCTTTCAC |
| plus 6 stop AS | GTGAAAGACGGGCTACACTTTCACCGC |
| plus 15 stop S | GGGTTTGCGGTGTTAGTGCAGCCCGTCTAACACCGTGCGGC |
| plus 15 stop AS | TGCCGCACGGTGTTAGACGGGCTGCACTAACACCGCAAACCC |
| UAA to AAA S | GGGTTTGCGGTGAAAGTGCAGCCCGTCTTTCACCGTGCGGC |
| UAA to AAA AS | TGCCGCACGGTGAAAGACGGGCTGCACTTTCACCGCAAACCC |
| UAA to UAU S | GGGTTTGCGGTGTATGTGCAGCCCGTCATACACCGTGCGGC |
| UAA to UAU AS | TGCCGCACGGTGTATGACGGGCTGCACATACACCGCAAACCC |
| LR to AA S | TGTAGTTGTGATCAAGCCGCCGAACCCATGCTTCAGTCAGCT |
| LR to AA AS | AGCTGACTGAAGCATGGGTTCGGCGGCTTGATCACAACTACA |
| Syn changes to attenuator S | CTCCGCGAACCCATGTTACAATCAGCAGATGCACAATCG |
| Syn changes to attenuator AS | CGATTGTGCATCTGCTGATTGTAACATGGGTTCGCGGAG |
